# A shared pathogen reservoir can tip widespread infection into mass mortality

**DOI:** 10.64898/2026.04.17.719273

**Authors:** Logan S. Billet, David K. Skelly, Erin L. Sauer

## Abstract

Pathogens that persist as low-intensity infections across many wildlife populations can drive mass mortality in others. Mass mortality is often abrupt, and the timing can be difficult to predict from host traits or environmental conditions alone. In a recent field study tracking ranavirus epizootics in wood frog (*Rana sylvatica*) breeding ponds, we found that environmental conditions and host features did not reliably distinguish die-off from non-die-off pond-years, whereas within-season trajectories of infection prevalence and environmental viral concentration did. Infection prevalence predicted subsequent environmental viral concentration throughout epizootics, whereas environmental viral concentration predicted subsequent infection prevalence and intensification primarily near die-off onset. This temporal asymmetry is consistent with a potential feedback operating through the shared viral environment. Here, we develop a compartmental model linking waterborne viral concentration to the rate at which low-intensity infections progress to high-intensity, high-shedding states within already-infected hosts. We show that a dose-dependent progression model generates the two-phase epizootic trajectory observed in natural die-offs: prolonged low-intensity circulation followed by a rapid increase in high-intensity infection and mortality as environmental viral concentration rises. The model exhibits a sharp mortality threshold between low-intensity circulation and mass mortality, governed mainly by the high-to-low intensity shedding ratio, host density, and pond volume. Existing explanations for die-off variation emphasize individual-level susceptibility, but our model demonstrates that dose-dependent environmental feedback acting on established infections can generate the transition from low-intensity infection to mass mortality in a homogeneous host population, even though variation in host susceptibility and environmental conditions may modify its timing and severity. This mechanism may apply in any system where hosts share a bounded environment, pathogen dose influences disease severity, and pathogen shedding increases with infection intensity.

## Introduction

Across wildlife disease systems, the same pathogen that circulates at low prevalence or low infection intensity in many populations can drive mass mortality events in others (Fey et al. 2015). This transition from widespread, inapparent infection to mass mortality can be abrupt, with population-wide collapse occurring over days to weeks following prolonged periods of low-intensity circulation (Vredenburg et al. 2010, Paul-Pont et al. 2014, Hall et al. 2018, Kock et al. 2018). Much of the effort to explain this variation has focused on individual-level susceptibility, seeking to identify host traits or environmental conditions (e.g., body condition, temperature) that increase the probability that individuals develop more severe infections (Brearley et al. 2013, Hing et al. 2016, Rollins-Smith 2017). However, while variation in individual susceptibility can help explain which hosts become infected and to what severity, it has often proven insufficient for predicting which populations experience die-offs, when die-offs occur, and how severe die-offs become, even in populations where infection is widespread (Lafferty and Holt 2003, Briggs et al. 2010, Langwig et al. 2012, Brunner et al. 2025a). The abruptness of the transition to mass mortality in many systems suggests the involvement of population-level processes not fully captured by variation in individual susceptibility. One candidate for such a process is the exposure environment itself.

Across a range of host-pathogen systems, the quantity of pathogen encountered can determine both whether infection establishes and how severe it becomes (Warne et al. 2011, Paul-Pont et al. 2015, McKenney et al. 2016, Niederwerder et al. 2019, Lunn et al. 2019). For example, mortality of Pacific oysters (*Crassostrea gigas*) during OsHV-1 exposures decreases with increasing water renewal rates, suggesting that viral accumulation and cumulative exposure, not variation in contact rate or host density, can drive differences in mortality (Petton et al. 2015). Importantly, the pathogen dose an individual experiences is not necessarily fixed at the point of initial exposure. In systems where infected hosts shed pathogen into a shared environmental reservoir, the collective output of the infected population may shape the ongoing exposure dose experienced by each host. Thus, rising prevalence may increase the pathogen concentration in the environment, potentially accelerating progression toward higher-intensity infection and further shedding, and thereby generate a positive feedback operating at the population level.

Larval amphibian populations offer an ideal system for evaluating whether pathogen buildup in a shared environment can influence progression toward higher-intensity infections. These animals often inhabit small, hydrologically isolated pools in dense populations where shed pathogens can accumulate in a shared water column. Ranaviruses are among the most significant pathogenic threats to amphibians globally, capable of driving rapid die-off events that can result in complete population mortality (Gray et al. 2009, Wheelwright et al. 2014, Price et al. 2014). However, while infection is often widespread across amphibian communities, die-offs are sporadic and difficult to predict (Brunner et al. 2025b). Several features of this system suggest that environmental pathogen accumulation could play a significant role in die-off dynamics: ranavirus-induced mortality is dose-dependent (Brunner et al. 2005, Warne et al. 2011), the virus can persist in natural pond water for days to weeks (Nazir et al. 2012, Johnson and Brunner 2014), and larval amphibians appear to lack the adaptive immune memory that would otherwise mitigate effects of repeated exposure to the same pathogen (Rollins-Smith 1998, Andino et al. 2012). Initial exposure dose also influences the magnitude and timing of peak viral load in larval amphibians, consistent with the possibility that ongoing environmental exposure could alter the trajectory of infection through effects on viral replication and immune control (Mihaljevic et al. 2019). However, whether continued exposure alters the trajectory of an already established infection has not been tested directly.

In a recent field study, we surveyed 40 wood frog (Rana sylvatica) breeding ponds for ranavirus epizootics across three years and found that environmental conditions and host features did not reliably distinguish die-off from non-die-off pond-years. Instead, lagged environmental viral concentration and infection prevalence were the most strongly supported predictors of subsequent infection spread and intensification, and their within-season trajectories distinguished die-off from non-die-off pond-years (Billet et al. 2026). Tadpole density at first detection was associated with faster environmental viral accumulation. Infection prevalence predicted subsequent waterborne viral concentration throughout epizootics, whereas waterborne viral concentration predicted subsequent prevalence and intensification primarily near the transition to mass mortality. This temporal asymmetry is consistent with the hypothesized feedback in which viral accumulation in the shared water column contributes to progression toward higher-intensity infection and shedding. However, whether environmental viral accumulation initiates the transition to mass mortality cannot be resolved using observational data alone.

Evaluating the potential role of viral accumulation in die-off occurrence and timing requires a modeling framework that can represent environmental pathogen concentration as a driver of progression toward higher-intensity infection within already-infected hosts, not just transmission to new ones. Existing theoretical frameworks have linked environmental exposure and within-host processes, including through parasite burden in classic macroparasite models (Anderson and May 1978, May and Anderson 1978) and infection load in host-parasite integral projection models (Wilber et al. 2016, 2021). Similarly, Mideo et al. (2008) described a case in which environmental pathogen abundance alters exposure and, in turn, infection progression within already-infected hosts, arguing that this reciprocal feedback requires explicit nesting of within-and between-host dynamics (i.e., essential nesting of models). However, in compartmental models of environmentally transmitted microparasites, the environmental reservoir has generally been treated as a driver of transmission to susceptible hosts rather than of progression toward higher-intensity infection in already-infected hosts (e.g., Codeço 2001, Tien and Earn 2010, Brunner and Yarber 2018). It remains unclear whether incorporating environmental pathogen concentration as a driver of infection progression in already-infected hosts can generate the transition from low-intensity circulation to mass mortality observed empirically (Hall et al. 2018, Billet et al. 2026). Answering this question has relevance beyond the ranavirus system, and especially in aquatic systems and aquaculture settings in which hosts shed pathogen into a shared medium with limited dilution and dose-dependent severity (Petton et al. 2015, McKenney et al. 2016).

Here, we develop a compartmental model that links environmental viral concentration to the rate at which low-intensity infections progress to high-intensity, high-shedding states, creating a feedback between the shared environment and the progression of established infections. Grounded in the temporal relationships observed in Billet et al. (2026), we ask whether dose-dependent infection progression can reproduce both the transition from widespread low-intensity infection to mass mortality and the asymmetric temporal pattern observed in the field, in which infected hosts shed virus throughout epizootics but environmental virus predicts subsequent infection intensification primarily near the transition to a die-off. To identify which features of the proposed mechanism are necessary to reproduce these dynamics, we compare three model variants of increasing complexity: a simple environmental transmission model with a single infected class, a staged-infection model with constant infection progression, and the full dose-dependent progression model.

## Methods

### Model formulation

We formalized the feedback described above as a compartmental ordinary differential equation (ODE) model in which environmental viral concentration influences the rate at which low-intensity infections progress to the high-intensity, high-shedding infections observed during ranavirus die-offs. The model tracks five state variables in a closed system (i.e., no immigration, emigration, or recruitment during the larval period): susceptible hosts (*S*), low-intensity infected hosts (*I*_low_), high-intensity infected hosts (*I*_high_), waterborne virus concentration (*W*, copies/mL), and infectious carcasses (*D*) (Figure 1A). This structure adapts the SIWR (susceptible-infected-water-recovered) framework of Tien and Earn (2010) by dividing infection into low-and high-intensity states, motivated by the distinct infection-intensity modes observed in natural populations and differences in shedding rates at different stages of ranavirus infection (Peace et al. 2019, Billet et al. 2026).

**Figure 1.**
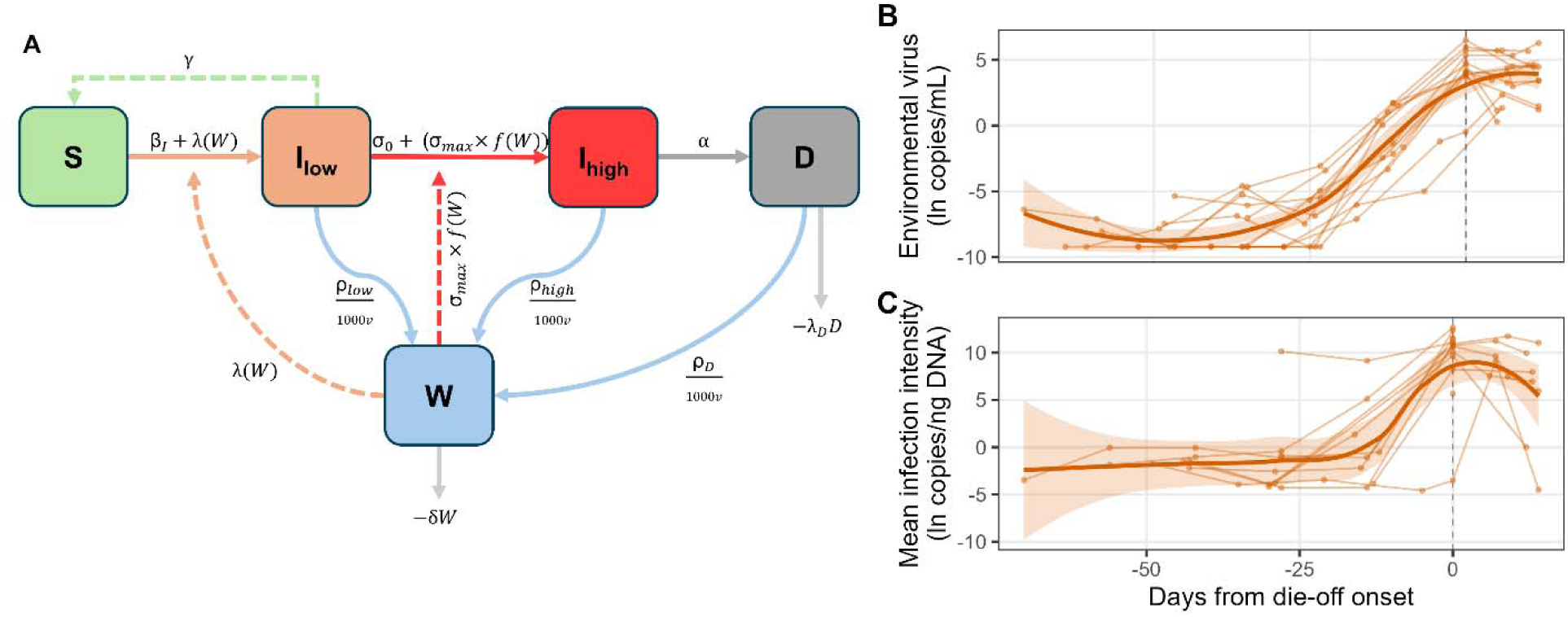
Model structure and empirical die-off trajectories. (A) Compartment flow diagram of the SIWR model with dose-dependent disease progression. Susceptible hosts (*S*) develop low-intensity infections (*I*_low_) through contact transmission (rate β*_I_*) and environmental force of infection, λ(*W*); low-intensity infections progress to high-intensity infections (*I*_high_) at rate σ(*W*), which depends on environmental viral concentration *W*. Hosts with high-intensity infections shed virus at rate ρ_high_, contributing ρ_high_/(1000*V)* to environmental viral concentration per host, and die at rate α, entering the carcass pool (*D*), which contributes virus into the environment at rate ρ*_D_*/(1000*V)*. The dashed green arrow indicates an optional recovery pathway (γ; set to zero in all baseline analyses). (B–C) Empirical trajectories from 13 die-off pond-years aligned to days from die-off onset (dashed vertical line at day 0): (B) environmental virus concentration (ln copies/mL) and (C) mean infection intensity among infected individuals (ln copies/ng DNA). Thin lines show individual pond-year trajectories; bold lines with shaded ribbons show loess smooths (span = 0.75) ± SE.

Hosts become infected through two routes: frequency-dependent contact transmission at rate β*_I_*(*I*_low_ + *I*_high_)/*N*, and environmental force of infection λ(*W*). For simplicity, we assume that direct-contact infectivity (β*_I_*) is the same for low-and high-intensity infections (but see Peace et al. 2019). This keeps the modeled difference between infection states confined to their environmental shedding rates and allows us to more clearly isolate the contribution of the environmental feedback. Ranavirus transmission does not cleanly follow density-or frequency-dependent formulations (Brunner et al. 2017), and our results are robust to either choice (see Results). The environmental force of infection follows a Hill-type dose-response (Appendix S1: Figure S11):

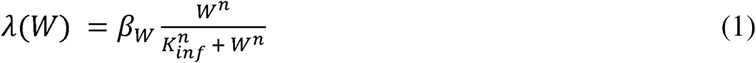

where β*_W_*is the maximum environmental force of infection, *K*_inf_ is the half-saturation concentration, and *n* is the Hill coefficient controlling the steepness of the dose-response relationship (Brouwer et al. 2017).

The central mechanism of the model is a dose-dependent progression function, σ(*W*), that determines the rate at which low-intensity infections transition to high-intensity infections. The rate at which existing infections intensify depends on the concentration of virus in the shared environment:

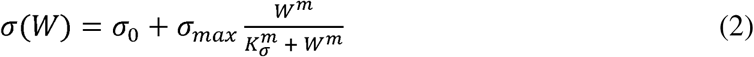

where σ_0_ is a baseline progression rate representing dose-independent progression, σ_max_is the maximum additional progression rate at saturating viral concentrations, *K*_σ_ is the half-saturation concentration for progression, and *m* is the Hill coefficient for the progression response. The Hill function acts as a flexible representation of nonlinear dose dependence, consistent with the dose-dependent mortality documented empirically in ranavirus and other amphibian-pathogen systems (Brunner et al. 2005, Briggs et al. 2010, Warne et al. 2011, Hall et al. 2020; Appendix S1: Figure S11).

Biologically, σ(*W*) is intended to act as a phenomenological representation of the possibility that ongoing environmental exposure alters the balance between viral replication and host immune control in an already-infected host, rather than as a model of a specific within-host mechanism. This interpretation is informed by within-host ranavirus models in which viral replication is opposed by a saturating immune response and initial dose alters both peak viral load and the timing of that peak (Mihaljevic et al. 2019). However, whether repeated exposure of an already-infected host can affect this balance and accelerate disease progression has not been tested directly, to our knowledge, and other biological mechanisms could produce a similar relationship between environmental exposure and progression. At the population level, this dose-dependent progression generates a positive feedback. High-intensity infected individuals shed virus at rates orders of magnitude higher than low-intensity infected individuals (ρ_high_ >> ρ_low_), increasing *W*, which in turn increases the progression rate of remaining low-intensity infections. When *W* is low and σ(*W*) ≈ 0, progression from low-to high-intensity infection is negligible, limiting the feedback between environmental viral concentration and high-shedding infections. As *W* increases toward and beyond *K*σ, σ(*W*) increases rapidly, increasing the rate of transition from low-to high-intensity infections and ultimately leading to mass mortality. The full system of ODEs is:

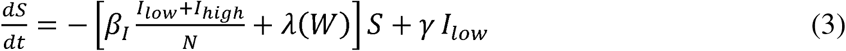

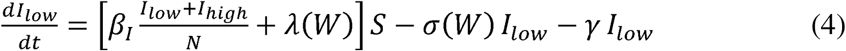

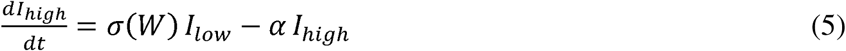

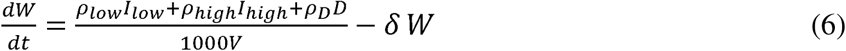

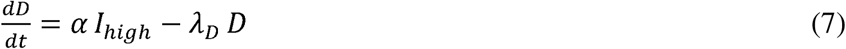

where α is the disease-induced mortality rate in high-intensity infections, γ is a low-intensity infection recovery rate (*I*_low_ to *S*), ρ_low_, ρ_high_, and ρ_D_ are viral shedding rates from hosts with low-intensity infections, high-intensity infections, and carcasses, respectively, *V* is pond volume, δ is the viral decay rate, and λ_D_ is the carcass removal rate. The recovery pathway γ is set to zero in the baseline parameterization and evaluated in sensitivity analysis.

### Model Comparison

We compared three model variants of increasing complexity: **(1)** the SIWR model (Model A)—a simple SIWR model with a single infected class and no dose-dependent feedback. This represents the structure of conventional compartmental models of environmentally transmitted microparasites, in which environmental pathogen concentration drives new infections but does not influence the progression of established infections; **(2)** the constant-progression model (Model B)—adds the *I*_low_/*I*_high_ staging and shedding asymmetry of the full model, but with progression fixed at σ_0_ + σ_max_/2 = 0.125 day^-1^ rather than depending on *W*. This represents a scenario in which infection progresses from low-to high-intensity states at a fixed rate, regardless of environmental context; **(3)** the dose-dependent model (Model C)—the complete model described by Eqs. 1-7, with dose-dependent progression σ(*W*). This captures the hypothesized feedback in which the rate at which existing infections progress from low-to high-intensity states depends on environmental viral concentration, linking population-level pathogen dynamics to progression among infection states. This is the focal model of the paper.

### Parameterization

We derived parameter values from a combination of published experimental, modeling, and field studies on ranavirus in amphibians, where available (Table 1); full derivations are provided in Appendix S1: Section S2. All concentration-dependent parameters in the model are expressed in genome-copy equivalents, because genome copies measured via qPCR-based environmental DNA (eDNA) are the field-measurable quantity in natural populations (Hall et al. 2016). Because qPCR-based eDNA quantifies viral genome copies rather than infectious virions, these concentrations should be interpreted as proxies for viral abundance rather than direct estimates of infectious dose (Appendix S1: Section S2). The model’s qualitative dynamics depend on ratios among concentration-dependent parameters rather than their absolute magnitudes (Appendix S1: Figure S9).

**Table 1.** Model parameters for baseline simulations.

| Parameter | Symbol | Baseline value | Units | Sensitivity range | Basis |
|---|---|---|---|---|---|
| Contact transmission rate | β<sub>*I*</sub> | 0.2 | day<sup>−1</sup> | 0.01–0.4 | C: Field buildup timing (range: Brunner and Yarber 2018, Peace et al. 2019) |
| Max environmental force of infection | β<sub>*W*</sub> | 0.2 | day<sup>−1</sup> | 0.01–0.5 | V: Set equal to β<sub>I</sub> |
| Half-saturation (infection) | *K*<sub>inf</sub> | 50 | copies/mL | 25–2,500 | D: Dose-response data (Warne et al. 2011) |
| Hill coefficient (Environmental transmission) | *n* | 2 | — | 1, 2 | V: Sigmoidal assumption |
| Hill coefficient (progression) | *m* | 2 | — | 2 | V: Sigmoidal assumption |
| Baseline progression rate | σ<sub>0</sub> | 0 | day<sup>−1</sup> | 0–0.10 | V: Field-constrained (Billet et al. 2026) |
| Max progression rate | σ<sub>max</sub> | 0.25 | day<sup>−1</sup> | 0–0.5 | D: Challenge study timescales (Pearman et al. 2004, Warne et al. 2011) |
| Half-saturation (progression) | *K*<sub>σ</sub> | 25 | copies/mL | 2.5–250 | C: Field die-off timing |
| Disease-induced mortality | α | 1/7 | day<sup>−1</sup> | 1/21–1/7 | D: (Brunner and Yarber 2018, Peace et al. 2019) |
| Low-intensity shedding | ρ<sub>low</sub> | 50 | copies/day | 50–5,000 | D: Titer-shedding scaling (Hall et al. 2020, Billet et al. 2026) at field low-intensity loads |
| High-intensity shedding | ρ<sub>high</sub> | 500,000 | copies/day | 50,000–500,000,000 | D: Informed by Hall et al. (2020) |
| Carcass leaching | ρ<sub>*D*</sub> | 500,000 | copies/day | 0, 500,000 | C/V: Set equal to ρ<sub>high</sub> (Brunner and Collins 2009, Peace et al. 2019) |
| Viral decay rate | δ | 0.25 | day<sup>−1</sup> | 0.03–0.5 | D: Johnson and Brunner (2014), Peace et al. (2019) |
| Carcass removal rate | λ<sub>*D*</sub> | 0.7 | day<sup>−1</sup> | — | E: Harp and Petranka (2006), Le Sage et al. (2019) |
| Pond volume | *V* | 10,000 | L | 1,000–50,000 | C: Set with N<sub>0</sub> to match field density |
| Initial population | *N*<sub>0</sub> | 20,000 | individuals | 2,000–100,000 | C: Set with *V* to match field density |
| low-intensity recovery rate | *γ* | 0 | day<sup>−1</sup> | 0–0.2 | V: Structural exploration |

Footnote: Basis codes indicate the evidential relationship between the cited work and the adopted value: E, value directly from published measurement; D, derived from empirical data; C, calibrated to field or model behavior; V, modeling convention or assumption. Sensitivity ranges report the values used in one-at-a-time perturbation analysis (Appendix S1: Figure S2).

The baseline progression rate is set to σ_0_ = 0, so the baseline model includes no dose-independent progression from low-to high-intensity infection. This assumption is consistent with our empirical observation that individuals with high tissue viral loads were largely confined to ongoing die-off events and were rare in ponds where environmental viral concentrations remained negligible (Billet et al. 2026). Though this strict assumption likely overstates the dependence of infection progression on environmental exposure, we use it to test whether the environmental pathway alone is sufficient to generate the observed die-off dynamics. The high-to-low shedding ratio we use (ρ_high_/ρ_low_ = 10^4^) is below the ∼10^5.5^ tissue-load asymmetry between low-and high-intensity infections in our field populations (Billet et al. 2026).

At baseline, the simulation uses *N*_0_ = 20,000 hosts in a pond of volume *V* = 10,000 L. This corresponds to a density of 2 individuals/L, comfortably within the range of field estimates for wood frog populations in vernal pools (Biesterfeldt et al. 1993, Harp and Petranka 2006). Viral concentration scales with the host-to-volume ratio rather than with population size or volume independently, and so the specific values of *N*_0_ and *V* matter mainly through their ratio. All simulations are seeded with 200 low-intensity infected hosts (*I*_low_(0) = 200; *S*(0) = *N*_0_ − 200; all other compartments at zero).

### Analytical and numerical methods

We evaluated the model by comparing the dynamics produced by each model variant qualitatively to the field patterns documented in Billet et al. (2026), instead of fitting directly to empirical time series. Our model is intended to test whether the hypothesized feedback mechanism can reproduce the two-phase die-off dynamics and asymmetric temporal pattern observed in natural populations, not to reproduce the trajectory of any individual pond. A primary outcome metric is cumulative mortality, calculated at time *t* as 1 − *N*_alive_(*t*)/*N*_0_, where *N*_alive_ = *S* + *I*_low_ + *I*_high_. Endpoint mortality is evaluated at *t*_final_ = 90 days, the approximate duration of the larval period. A die-off is defined as cumulative mortality exceeding 50% of the initial population.

We computed the basic reproduction number *R*_0_ using the next-generation matrix (NGM) method (van den Driessche and Watmough 2002); the full NGM derivation is provided in Appendix S1: Section S1. To identify the parameter conditions under which the system transitions between low-intensity infection and mass mortality, we varied the shedding ratio ρ_high_/ρ_low_ across eight orders of magnitude (10^0^ to 10^8^) by varying ρ_high_ while ρ_low_ was held fixed, and recorded cumulative mortality. We extended this to two dimensions by jointly varying shedding ratio and pond volume (*V* from 1,000 to 100,000 L) at fixed host abundance (*N*_0_ = 20,000, effectively varying host density from 0.2 to 20 individuals/L).

To evaluate robustness to parameter uncertainty, we combined one-at-a-time (OAT) perturbation over the ranges in Table 1 with targeted two-dimensional sweeps of the parameters that were the least empirically constrained (Appendix S1: Figures S6-S9) and tested structural robustness by varying the Hill coefficient for environmental transmission (n = 1 vs. 2) and the functional form of the progression function (Appendix S1: Section S3).

All analyses were conducted in R version 4.3.2 (R Core Team 2023). All deterministic simulations used the lsoda solver in the R package deSolve (Soetaert et al. 2010).

## Results

Under our baseline parameterization, the dose-dependent model (Model C) produces a two-phase epizootic consistent with patterns observed in naturally occurring ranavirus die-offs (Figure 1B-C; Figure 2). During the first ∼37 days, mortality and environmental viral concentration remain low. Contact transmission (β*_I_*) results in a slowly building low-intensity infection pool, but environmental viral concentration remains low and both environmental transmission and dose-dependent progression are negligible. Each transition to high-intensity infection, however, increases per-host shedding by four orders of magnitude, and as high-intensity infection prevalence reaches approximately 5% of the population, cumulative shedding pushes *W* above the progression half-saturation concentration *K*_σ_. This, in turn, produces a rapid shift, wherein high-intensity infection prevalence peaks within a week of W passing *K*_σ_, peak environmental viral concentration reaches ∼1,800 copies/mL, and cumulative mortality exceeds 90% by day 60.

**Figure 2.**
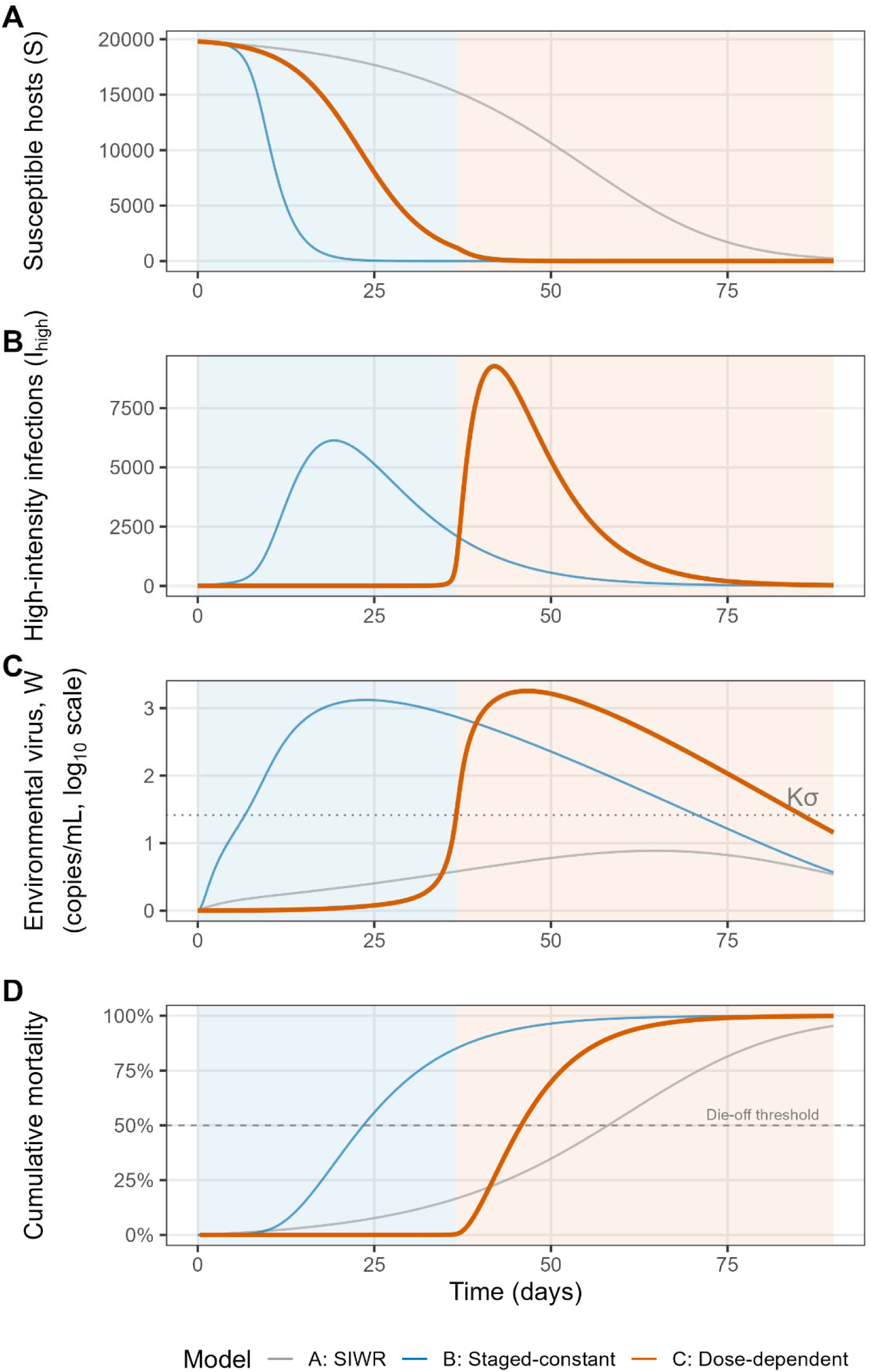
Comparative dynamics of three model variants. Time series of (A) susceptible hosts, (B) high-intensity infections, (C) environmental virus concentration (log_10_ scale, with *K*_σ_ = 25 copies/mL marked), and (D) cumulative mortality for Model A (SIWR, grey), Model B (staged-constant, blue), and Model C (dose-dependent, orange). Phase shading indicates the early buildup phase (blue) and rapid progression phase (orange), defined by the time at which *W* exceeds *K*_σ_ in Model C. The dashed line in (D) marks the 50% die-off threshold. All three models produce high mortality, but Model C exhibits a distinctive biphasic trajectory: an extended buildup phase with negligible high-intensity infection and mortality, followed by rapid increases in high-intensity infection, environmental virus concentration, and mortality.

All three model variants produce high cumulative mortality (>90%) with baseline parameters but differ substantially in the temporal dynamics of mortality (Figure 2, Appendix S1: Figure S1, Table S1). Model A (single infected class, no staging) produces a relatively smooth mortality curve with no distinct acceleration phase, reaching 50% mortality at day ∼58. Model B (staged infection with constant progression) generates faster dynamics because the shedding asymmetry between low-and high-intensity infections increases environmental viral concentration as high-intensity infections accumulate. However, because progression is fixed, transitions to high-intensity infections begin immediately following infection and occur gradually, with 50% mortality occurring at day ∼24. Only Model C (dose-dependent progression) reproduces a two-phase pattern characterized by a prolonged period of near-zero mortality followed by an abrupt increase in mortality as environmental viral concentration reaches levels at which σ(*W*) increases rapidly, reaching 50% mortality at day ∼46. The qualitative distinction between models therefore lies mainly in the timing and sharpness of the increase in mortality rather than in cumulative mortality at the end of the 90-day simulation.

Threshold analysis across the shedding ratio ρ_high_/ρ_low_ shows a sharp mortality threshold in Model C near a shedding asymmetry of approximately 1,000-fold (log_10_ ratio ≈ 3.0; Fig 3A). Cumulative mortality increases from near-zero to near-total over approximately 0.2 log_10_ units. This is because below this threshold, environmental viral concentration remains below *K*σ, limiting progression to high-intensity infection. above that threshold, *W* reaches concentrations at which σ(*W*) increases quickly, leading to rapid accumulation of high-intensity infections and mass mortality. Neither Model A nor Model B exhibits a comparable threshold. The threshold shedding ratio in Model C (10^3^) is below the baseline parameterization (10^4^) and the field tissue-load asymmetry (∼10^5.5^; Billet et al. 2026), suggesting that the shedding asymmetry required to generate the sharp mortality threshold falls within an empirically plausible range.

**Figure 3.**
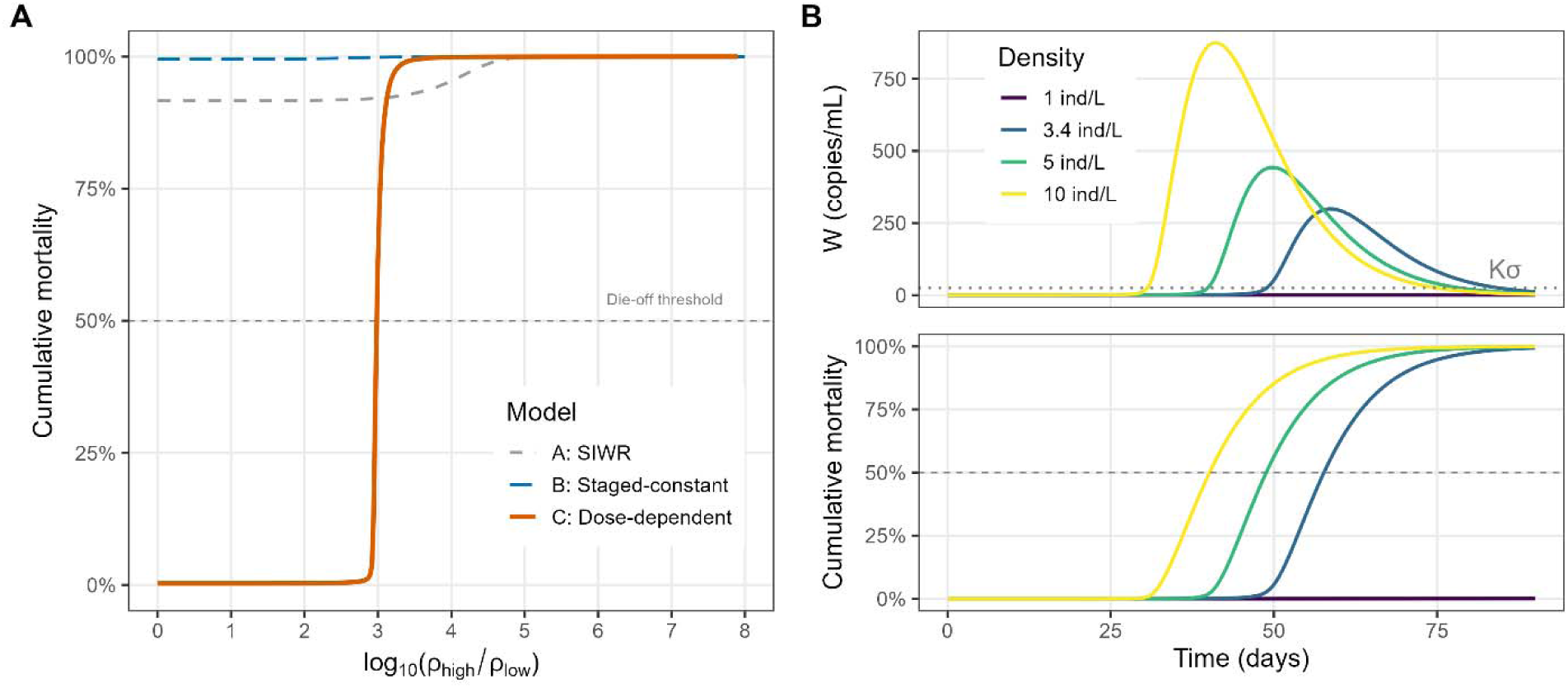
Sharp mortality threshold and density-dependent dynamics in Model C. (A) Threshold analysis showing cumulative mortality as a function of the high-to-low shedding ratio (log_10_(ρ_high_/ρ_low_)) for all three models. Model C (orange) exhibits a sharp mortality threshold spanning ∼0.2 log units, while Model B (blue, long-dashed) produces uniformly high mortality regardless of shedding ratio and Model A (grey, dashed) shows a gradual, monotonic increase. Dashed horizontal line marks the 50% die-off threshold. (B) Density-dependent dynamics at near-threshold shedding (ρ_high_= 50,000 copies/day): environmental virus trajectories (top, with *K*_σ_ marked) and cumulative mortality (bottom) at four host densities (range: 1–10 ind/L), achieved by varying *N*_0_ at constant volume. Higher densities produce greater aggregate shedding per unit volume, raising *W*, increasing σ(*W*), and producing rapid increases in high-intensity infections and mortality.

A joint sweep of shedding ratio and pond volume at fixed *N*_0_ = 20,000 shows that the shedding ratio required for a die-off increases with pond volume, reflecting dilution of shed virus across larger water volumes (Appendix S1: Figure S12). At *V* = 1,000 L (20 individuals/L), the 50% mortality boundary occurs at a high-to-low shedding ratio of approximately 10^1^, whereas at *V* = 10,000 L (baseline; 2 individuals/L), the 50% mortality boundary occurs near a shedding ratio of ∼10^3^ and rises to ∼10^5^ at *V* = 100,000 L (0.2 individuals/L). *R*_0_ at baseline parameters is 1.4 for Model A and 3.0 for Model B. For Model C, *R*_0_ is undefined because low-intensity infected hosts have no exit pathway when σ_0_ = 0 and γ = 0, making the infected-state transition matrix singular (Appendix S1: Section S1). However, the initial epizootic growth rate shows that infection can invade under baseline parameterization. For Models A and B, *R*_0_ is independent of pond volume under frequency-dependent contact transmission, whereas the initial epizootic growth rate provides the corresponding invasion metric for Model C. These invasion metrics do not predict the volume-dependent die-off boundary because they describe dynamics near the disease-free state, whereas the mortality threshold depends on whether environmental viral concentration reaches levels at which σ(*W*) increases rapidly, which depends on host density and pond volume (Appendix S1: Table S1).

At the baseline high-intensity shedding rate (ρ_high_ = 500,000), the 50% mortality boundary occurs at a pond volume of approximately 31,000 L (density ≈ 0.64 individuals/L). At near-threshold shedding (ρ_high_ = 50,000 copies/day; ρ_high_/ ρ_low_ = 10^3^), the boundary occurs at approximately 10,900 L (density ≈ 1.84 individuals/L; Appendix S1: Figure S12). Host density produces the complementary effect, where at fixed volume and near-threshold shedding, increasing host density raises aggregate shedding per unit volume, pushing *W* toward *K*_σ_. At 1 ind/L, environmental viral concentration remains below *K*_σ_ throughout the simulation, whereas at 3.4 ind/L and above, *W* crosses *K*_σ_ and dose-dependent progression increases rapidly (Figure 3B). This density dependence aligns with the empirical observation that tadpole density at first detection predicted the rate of environmental viral accumulation across field populations (Billet et al. 2026). In the model, higher host densities increase aggregate shedding per unit volume, elevating *W* more rapidly and reducing the time required for σ(*W*) to reach high values.

Adding recovery from low-intensity infection (γ > 0) introduces competition between two outcomes for low-intensity infected hosts: clearance (at rate γ) and progression to high-intensity infection (at rate σ(*W*)). When environmental viral concentration is low, recovery can remove infections before progression occurs; as *W* increases, σ(*W*) increases and infection progression can outpace recovery, leading to a rapid accumulation of high-intensity infections. At γ = 0.05 day^-1^ (approximately 3-week mean clearance time), Model C still produces die-offs exceeding 90% mortality when environmental viral accumulation is sufficient to generate rapid progression, but parameter combinations that produce moderate mortality (50-80%) in the baseline model (γ = 0) instead result in outcomes with negligible mortality (Appendix S1: Figure S5). The recovery pathway thus sharpens the distinction between die-off and non-die-off outcomes without eliminating die-offs. Because recovery from low-intensity ranavirus infection remains poorly documented, this analysis points toward recovery as an important source of uncertainty.

One-at-a-time sensitivity analysis identifies six parameters for which variation across the tested range shifts cumulative mortality from near-zero to near-total mortality (Appendix S1: Figure S2): maximum progression rate (σ_max_), contact transmission rate (β*_I_*), progression half-saturation (*K*_σ_), pond volume (*V*), low-intensity infection recovery rate (γ), and initial population size (*N*_0_). All six share the property that perturbation can move the system across the die-off threshold, consistent with the sharp mortality threshold identified in the shedding-ratio analysis. In contrast, the low-intensity shedding rate (ρ_low_) and disease-induced mortality rate (α) have little individual influence on cumulative mortality (OAT mortality range < 0.094). Targeted two-dimensional sweeps confirm that the model’s qualitative behavior is robust to joint uncertainty in the parameters with the least direct empirical support: die-off dynamics persist across 56% of the *K*_inf_ × *K* parameter combinations tested (Appendix S1: Figure S6), across δ = 0.1-0.6 day^-1^(Appendix S1: Figure S7), and under proportional rescaling of all concentration-dependent parameters across four orders of magnitude (Appendix S1: Figure S9). The two-phase die-off pattern is also robust to the alternative progression functions tested in Appendix S1: Section S3, removal of carcass contributions, and density-dependent contact transmission (Appendix S1: Figure S10). The exception is a step-function formulation, which fails because low-intensity shedding alone cannot raise *W* to *K*_σ_ (see Appendix S1: Section S3).

## Discussion

Ranavirus epizootics are frequently characterized by widespread, low-intensity infection followed by an abrupt transition to a die-off. Our recent observational work found that infection prevalence predicted subsequent waterborne viral concentration throughout epizootics, whereas waterborne viral concentration predicted subsequent infection prevalence and intensification primarily when die-off-onset visits were included (Billet et al. 2026). In the present study, we show that a compartmental model linking environmental viral concentration to the rate at which low-intensity infections progress to high-intensity infections qualitatively reproduces the rapid transition from widespread, low-intensity infection to mass mortality observed in naturally occurring epizootics. The inclusion of dose-dependent progression produces this asymmetry. When environmental viral concentration is low, progression to high-intensity infection is negligible, but as the low-intensity infected pool grows and cumulative shedding elevates concentration, individuals begin to develop high-intensity infections, eventually raising environmental viral concentrations into the range where σ(*W*) increases rapidly and further increases progression and shedding. Importantly, the shift from low-intensity infection to mass mortality occurs in the model without requiring temporal changes in host susceptibility, showing that environmental viral accumulation may be sufficient to generate this pattern even though host and environmental variation may modify its timing and severity.

Several modeling frameworks already link environmental exposure or parasite burden to changes in infection severity, including classic macroparasite models and host-parasite integral projection models (Anderson and May 1978, May and Anderson 1978, Wilber et al. 2016, 2021). However, compartmental models of environmentally transmitted microparasites have generally connected environmental pathogen concentration to the force of infection rather than the progression of established infections (Codeço 2001, Tien and Earn 2010). This limitation has long been recognized: Codeço (2001) explicitly noted the assumption that inoculum size affects infection probability but not severity, and subsequent work has called for models that extend the influence of dose to disease severity (Grad et al. 2012, Garira et al. 2014). Even dedicated treatments of dose-response in environmental transmission models have focused primarily on the probability of infection rather than infection severity or progression (Brouwer et al. 2017, Lanzas et al. 2020). Related microparasite models have linked environmental pathogen concentration to continuous within-host infection load (Feng et al. 2013) or represented disease progression through pathogen-load dynamics and load-dependent mortality (Briggs et al. 2010). Our model builds on these precedents by allowing environmental pathogen concentration to alter transitions between discrete infection states within already-infected hosts, providing a rough population-level approximation of the reciprocal exposure-progression coupling that Mideo et al. (2008) argued would require explicit nesting of within-and between-host dynamics. We use two infection states as a simplified representation of low-and high-intensity infection modes observed empirically (Billet et al. 2026), which allows us to better isolate the environmental feedback without adding complexity unnecessary to test the mechanism. This simplification represents progression as an immediate shift in individual shedding, and additional infection stages could capture more gradual changes in pathogen load and shedding (e.g., Peace et al. 2019).

Similar threshold behavior has been described in models with nonlinear or reinforcing feedbacks on transmission, such as dose-dependent and environmentally mediated transmission models. These nonlinearities can generate invasion thresholds, bistability, and hysteresis, such that *R*_0_ may not always predict whether a pathogen can invade or persist (Regoes et al. 2002, Roberts 2007, Joh et al. 2009, Brouwer et al. 2017). Our model has a similar nonlinearity in the environmental force of infection, λ(*W*). At the baseline Hill coefficient (n = 2), the environmental force of infection changes little when *W* is near zero, so the earliest invasion dynamics are driven primarily by contact transmission (Appendix S1: Section S1). However, this nonlinearity in environmental force of infection does not explain the sharp mortality threshold observed in Model C; Model A uses the same λ(*W*) function but shows a relatively gradual increase in mortality as the shedding ratio increases (Figure 3A). When n = 1, environmental force of infection responds more strongly at low viral concentrations, but the sharp mortality threshold remains essentially unchanged (Appendix S1: Figures S3-S4). These comparisons show that the threshold depends on the dose-dependent progression pathway acting on established infections rather than on the nonlinear environmental force of infection. Our results complement earlier work on transmission-side environmental feedbacks by showing that nonlinear responses in established infections can also generate abrupt shifts in population-level mortality.

The sharp mortality threshold generated by dose-dependent progression from low-to high-intensity infection raises an important biological question: why should additional exposure alter the course of an infection that is already established? Traditionally, models of microparasite infections assume that it should not. Within-host replication is expected to overwhelm any subsequent inoculum rapidly enough that infection outcomes are treated as functionally independent of additional exposures once infection is established (Anderson and May 1979). The dose-dependent progression function that we describe, σ(*W*), is a departure from this assumption. The population-level dynamics we describe do not depend on a specific within-host process by which continued exposure may alter infection progression, but identifying plausible mechanisms is important if the feedback is to be tested experimentally. One possibility is that within-host processes that are sensitive to the initial exposure dose may remain responsive to additional exposure after infection is established. The initial exposure dose can alter both the magnitude and timing of peak viral load in larval amphibians (Mihaljevic et al. 2019), suggesting that viral replication and immune control may be sensitive to the amount of virus encountered by the host. A recent model integrating dose dependence into transmission dynamics captures this effect by allowing the environmental pathogen concentration at the time of infection to determine the infection class that hosts enter, while subsequent progression is independent of continued environmental exposure (Chowdhury et al. 2026). In contrast, σ(*W*) allows environmental pathogen concentration to continue altering the rate of progression between infection states after infection has already been established. A second possibility is that some additional inoculum may escape host immune control and directly contribute to within-host viral replication after infection is already established. Tadpoles appear to mount a delayed and limited innate immune response, and ranaviruses have multiple mechanisms of immune evasion (Grayfer et al. 2012, 2015, Andino et al. 2012). Infection-induced damage to epithelial barriers could further increase opportunities for additional virus to enter and replicate in tissues not yet reached by the initial infection (Forzán et al. 2017). Third, infection-induced declines in host body condition could progressively reduce resistance to further exposure, creating a ‘vicious circle’ in which infection increases susceptibility to more severe infection (Beldomenico and Begon 2010). However, direct experimental evidence that continued pathogen exposure alters infection progression after infection has already been established is currently lacking. A useful test could expose hosts to a common low initial dose and then vary the duration of continued low-level environmental exposure to evaluate whether prolonged exposure influences viral load, disease progression, and mortality.

Existing explanations of ranavirus die-off timing have primarily emphasized individual-level processes, including ontogenetic changes in susceptibility and environmental conditions such as temperature. Brunner et al. (2015), for example, modeled stage-dependent susceptibility increasing toward metamorphosis (Warne et al. 2011) as the primary driver of summer die-off timing. However, die-offs observed across our field populations began across a wide temporal window (May 24-July 12), corresponding to wide ranges of mean developmental stages (Gosner 28.8-41.2) and mean temperatures over the preceding 7 days (12.2-20.2°C; Billet et al. 2026). These observations suggest that ontogeny and temperature alone do not determine die-off timing, but they do not imply they are unimportant. Our model highlights environmental viral accumulation as a population-level process that can interact with these other drivers. Developmental stage, temperature, and individual variation in susceptibility, for example, could alter rates of infection, infection progression, shedding, and viral persistence, thus modifying the conditions under which the feedback might result in mass mortality. Within our model, recovery acts as one example of this kind of modification. When low-intensity infections can be cleared, recovery can compete with dose-dependent progression and make intermediate outcomes less common, so populations are more likely to either recover with limited mortality or experience a die-off (Appendix S1: Figure S5).

Die-off timing in the model emerges from the rate at which environmental virus accumulates through collective shedding, a process influenced by transmission rates, host density, and pond volume. Density in the model does not predict die-off occurrence directly, but rather modulates the rate of environmental viral accumulation, which determines whether the viral concentration reaches levels at which σ(*W*) increases rapidly within the larval period. This is consistent with the empirical association between tadpole density at first detection and faster eDNA accumulation; density did not directly distinguish die-off from non-die-off pond-years (Billet et al. 2026). The environmental feedback mechanism also informs a specific prediction about experimental outcomes. In small experimental venues, the host-to-volume ratio is likely high enough that environmental viral concentration reaches these levels rapidly regardless of treatment, which may explain why experimental manipulations of stressors that influence individual susceptibility have not scaled to population-level outcomes (Haislip et al. 2012, Reeve et al. 2013, Brunner et al. 2025a). More broadly, real populations contain individual variation in susceptibility that our model omits. Including such heterogeneity could allow the most susceptible individuals to progress to high-shedding states earlier, accelerating environmental viral accumulation and potentially shifting the mortality threshold relative to our homogeneous model.

The qualitative dynamics of the model are robust to relaxing the assumption that baseline progression (σ_0_) is zero so long as σ_0_remains small relative to σ_max_. At low baseline progression rates, low-intensity shedding alone is insufficient to raise environmental viral concentrations into the range where σ(*W*) increases rapidly, and the temporal asymmetry between low-intensity circulation and mass mortality persists. The model also omits acquired immunity, host heterogeneity, and spatial structure, all of which likely have some relevance in other host-pathogen systems, more diverse amphibian assemblages, and larger bodies of water. The closed-population assumption is realistic for wood frog tadpoles confined to vernal pools during the larval period, but should be relaxed for systems with continuous recruitment or host movement between water bodies. Additional and more heterogeneous exposure routes exist beyond the waterborne pathway modeled here, including direct contact with low-and high-intensity infected conspecifics and necrophagy of infectious carcasses. However, the core mechanism, continued pathogen exposure altering infection progression within already-infected hosts, could in theory operate through any of these pathways. Future models could extend the exposure-dependent progression mechanism to incorporate these additional transmission pathways more explicitly.

The feedback our model demonstrates has the potential to operate wherever hosts share a physically bounded environment, pathogen dose influences disease severity in already-infected hosts, and shedding increases with disease severity. A mechanistic parallel comes from amphibian chytridiomycosis, where Vredenburg et al. (2010) observed widespread low-intensity *Batrachochytrium dendrobatidis* (Bd) infection followed by rapid increases in infection intensity past a lethal infection threshold, and Briggs et al. (2010) showed that collective zoospore accumulation in the shared aquatic environment can determine whether Bd infections progress to lethal loads. In aquaculture and fish disease systems, comparable patterns have been documented. These include mortality that scales with cumulative viral shedding rather than initial challenge dose, density-dependent acceleration of mortality accompanied by proportional increases in waterborne viral concentration, and higher mortality among hosts sharing water than among isolated individuals at identical doses and full infection prevalence (Hershberger et al. 2010, Petton et al. 2015, McKenney et al. 2016, Kim et al. 2025). These outcomes have generally been attributed to host-level processes such as immune suppression or elevated contact rates under crowding, but the environmental feedback we describe here provides a complementary population-level mechanism consistent with these patterns.

Our model shows that allowing environmental pathogen concentration to influence infection progression within established infections is sufficient to produce an abrupt shift from widespread, low-intensity infection to mass mortality. Within this framework, the parameters governing environmental pathogen accumulation (host density, shedding asymmetry, the volume of the shared environment) determine whether and when die-offs occur. Because environmental pathogen abundance can be tracked in the field using eDNA and, increasingly, eRNA as measurable proxies (Farrell et al. 2021, Bass et al. 2023), monitoring the exposure environment alongside infection prevalence may help identify populations at risk for die-offs, with similar feedbacks potentially operating wherever hosts contribute pathogens to a shared, bounded environment.

## Acknowledgments

We thank Jason Hoverman, Vanessa Ezenwa, Brandon Ogbunu, and Trenton Garner for useful suggestions and conversations. We thank Adriana Rubinstein, L. Kealoha Freidenburg, Carolyn Skotz, Yara Alshwairikh, Sydney Nelson, and Karinne Tennenbaum for their assistance with fieldwork and/or labwork for the study motivating this work. We acknowledge the startup support provided to E.L.S. from the School of Environmental and Biological Sciences at Rutgers University. The field observations motivating this work were supported by funds awarded to L.S.B from the Kohlberg-Donohoe Research Fellowship, the Yale Institute for Biospheric Studies, the Yale Law School LEAP Student Grant Program, the AMNH Theodore Roosevelt Memorial Fund, and Sigma Xi Grants in Aid of Research, and provided to D.K.S from Yale University.

## Conflict of Interest Statement

The authors declare no conflicts of interest.

## Notes

### Competing Interest Statement

The authors have declared no competing interest.

### Summary of Updates

In response to the constructive comments from two reviewers, we have substantially revised the manuscript. In particular, we expanded the Introduction and Discussion to better position our work within existing theory on environmentally transmitted pathogens, nonlinear transmission dynamics, and feedbacks between exposure and infection processes. We also expanded our consideration of biological mechanisms that could plausibly produce the exposure-progression relationship represented in the model and revised terminology and framing throughout in response to both reviewers, including replacing subclinical and clinical infection with low-intensity and high-intensity infection.

