## Appendix 1 for "A shared pathogen reservoir can tip widespread infection into mass mortality"

### Section S1: *R*_0_ Derivation (Next-Generation Matrix)

Because the carcass compartment *D* reaches pseudo-equilibrium (*D** = α·*I*_high_/λ_D_) on a timescale much shorter than the epizootic dynamics, we reduced the disease subsystem to three compartments (*I*_low_, *I*_high_, *W*) and incorporate carcass-mediated shedding as an effective addition to the high-intensity shedding term. Following van den Driessche and Watmough (2002), the transmission matrix **F** (new infections generated per unit time by each infected compartment) and the transition matrix **V** (rates of transition out of each compartment, minus inter-compartment transfers) are linearized at the disease-free equilibrium (*S*, *I*_low_, *I*_high_, *W*, *D*) = (*N*_0_, 0, 0, 0, 0).

**Transmission matrix F** (3 × 3):

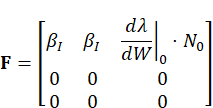

where dλ/d*W*|_0_ denotes the derivative of the environmental force of infection evaluated at *W* = 0. For the Hill-type dose-response λ(*W*) = β*_W_* · *W^n^*/(K*^n^*_inf_ + *W_n_*), the general derivative is *n*·β*_W_*·*W*^(n−1)^·*K^n^*_inf_ / (*K^n^*_inf_ + *W^n^*)^2^, which evaluates at W = 0 to:

*n* = 1: dλ/d*W*|_0_ = β*_W_* / K_inf_

*n* ≥ 2: dλ/d*W*|_0_ = 0

For all analyses using the baseline Hill coefficient *n* = 2, environmental transmission does not contribute to **F** at the disease-free equilibrium (DFE), and when **V** is nonsingular, *R*_0_ is governed entirely by the contact transmission pathway.

**Transition matrix V** (3 × 3):

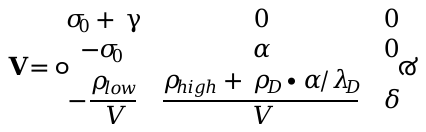

The term ρ*_D_*·α/λ*_D_* in the (*W*, *I*_high_) entry incorporates the steady-state carcass contribution: each high-intensity infected host that dies (at rate α) produces a carcass that sheds at rate ρ*_D_* until removed (at rate λ*_D_*), yielding an effective additional shedding rate of ρ*_D_*·α/λ*_D_* per high-intensity infection.

For Model C at baseline parameters (σ_0_ = 0, γ = 0), the first diagonal entry of **V** is zero, making **V** singular and *R*_0_ undefined. The singularity reflects the model's structural assumption that hosts with low-intensity infections have no exit pathway absent environmental feedback, not unbounded transmission. As σ_0_ and γ approach zero from positive values, however, the corresponding finite *R*_0_ increases without bound. For Model C, the biologically meaningful invasion metric is the initial epizootic growth rate (the dominant eigenvalue of **F** − **V**, equal to β*_I_* = 0.2 day^-1^ when environmental transmission is absent at the DFE), which confirms that infection can invade but does not predict outbreak severity. For Model B, σ_0_ is replaced by the constant progression rate σ_const_ = 0.125 day^-1^, yielding a non-singular **V** and finite *R*_0_ = 3.0. For Model A, the single-class structure reduces the system to a simple SIWR formulation with *R*_0_ = 1.4 (Table S1).

**Section S2: Parameter Justifications**

This section provides detailed derivations for each parameter in Table 1. All parameters are expressed in genome-copy equivalents. The experimental dose-response literature for ranavirus reports inoculation doses in plaque-forming units (PFU), whereas field-measurable concentrations are quantified as genome copies via qPCR-based environmental DNA (eDNA). Because qPCR-based eDNA quantifies viral genome copies rather than infectious virions, these concentrations should be interpreted as proxies for viral abundance rather than direct estimates of infectious dose. The genome-copy-to-PFU ratio for ranavirus is not clearly defined, and likely varies with viral strain, quantification method, etc. The absolute values of concentration-dependent parameters (shedding rates, half-saturation concentrations, viral decay) are therefore uncertain. However, the model's qualitative dynamics depend on ratios among these parameters rather than their absolute magnitudes: proportionally rescaling all concentration-dependent parameters (ρ_high_, ρ_low_, ρ_D_, *K*_σ_, and *K*_inf_) across four orders of magnitude leaves cumulative mortality and onset timing unchanged (Appendix S1: Figure S9). This rescaling invariance suggests that the unit-conversion uncertainty does not affect the qualitative conclusions.

**Transmission rates.** The contact transmission rate and maximum environmental force of infection (β*_I_* = β*_W_* = 0.2 day^-1^) were set equal, as we had no strong empirical basis for weighting one route over the other at baseline. This value falls within the range used in previous ranavirus models (Brunner and Yarber 2018, Peace et al. 2019) and produces buildup timing comparable with that observed in natural populations (Hall et al. 2018, Billet et al. 2026).

**Half-saturation concentration for infection.** The half-saturation concentration for infection (*K*_inf_ = 50 copies/mL) was informed by dose-response data, in which the LD_50_ for wood frog tadpoles in bath exposure was 10^2.37^ PFU/mL, and the ID_50_ was below the lowest dose tested (<10^1.4^ PFU/mL; Warne et al. 2011, Brunner et al. 2025), suggesting that infection can be established at very low exposure concentrations. Because *K*_inf_ represents a continuous exposure process rather than a single acute bath challenge, cumulative exposure over days at moderate concentrations can deliver an effective dose exceeding that of a brief high-concentration pulse.

**Hill coefficients.** We use Hill coefficients of *n* = 2 for environmental transmission and *m* = 2 for progression, producing sigmoidal responses consistent with the nonlinear dose-mortality relationships observed in experimental challenge studies (Brunner et al. 2005, Warne et al. 2011) and the nonlinear temporal pattern observed in field populations (Billet et al. 2026). We evaluated structural robustness to the environmental-transmission Hill coefficient by comparing *n* = 1 and *n* = 2 while holding *m* = 2.

**Progression half-saturation.** The progression half-saturation (*K*_σ_ = 25 copies/mL) was selected to produce progression dynamics consistent with the rapid transition from low- to high-intensity infections observed in field die-offs (Hall et al. 2018, Billet et al. 2026). *K*_σ_ is the environmental viral concentration at which the exposure-dependent component of progression reaches half its maximum rate. We interpret this parameter phenomenologically rather than as a directly measured biological threshold. Within-host ranavirus models in which viral replication is opposed by a saturating immune response and initial dose alters both peak viral load and the timing of the peak provide biological plausibility for dose-sensitive infection progression (Mihaljevic et al. 2019), although this remains to be rigorously tested. The maximum infection progression rate (σ_max_ = 0.25 day^-1^) was informed by experimental challenge studies that provide timescales for the development of high-intensity infections under high-dose conditions (Pearman et al. 2004, Warne et al. 2011), and corresponds to a progression timescale of approximately 4 days at saturation.

**Disease-induced mortality.** For the rate of disease-induced mortality, we selected a value consistent with available evidence, α = 1/7 day^-1^. This corresponds to a 7-day mean time from high-intensity infection to death, and falls between the 18-day value used by Brunner and Yarber (2018) in their transmission model (which encompasses the full infectious period) and the ~5.1-day estimate from Peace et al. (2019), which was derived from experimental time-to-death data encompassing both latent and infectious stages.

**Shedding rates.** The low-intensity shedding rate (ρ_low_ = 50 copies day⁻¹) was selected with reference to Hall et al. (2020), who found that shed eDNA concentrations scale ~1:1 with gastrointestinal tissue viral titers on the log scale (β = 0.95). The low-intensity component of the infection-intensity distribution in our field populations had a mean of approximately 0.08 copies/ng DNA (Billet et al. 2026), supporting a low shedding rate relative to hosts with high-intensity infections. The high-intensity shedding rate (ρ_high_ = 500,000 copies day⁻¹) was informed by Hall et al.'s (2020) measurements from high shedders at 6 days post-inoculation (up to ~227,000 copies day^-1^), and is relatively conservative relative to terminal-stage shedding if extrapolated from samples collected during die-offs. The resulting shedding ratio ρ_high_/ρ_low_ = 10^4^ is below the ~10^5.5^ difference between low- and high-intensity components of the tissue viral-load distribution in our field populations (Billet et al. 2026). Carcass leaching (ρ_D_ = 500,000 copies day^-1^) follows the convention of Peace et al. (2019), setting carcass viral output equal to the high-intensity shedding rate, which is consistent with the high titers documented in carcasses (Brunner and Collins 2009).

**Viral decay.** The viral decay rate (δ = 0.25 day^-1^) follows Peace et al. (2019), based on Johnson and Brunner's (2014) data on ranavirus persistence in natural pond water. Culturable virus in unmanipulated pond water declines rapidly (T90 < 1 day), whereas viral DNA persists for much longer. The value δ = 0.25 day^-1^ (T90 ≈ 9 days) corresponds approximately to the persistence of virus under filter-sterilized conditions in the same study, and we interpret *W* as an effective proxy for environmental viral abundance that declines faster than total viral DNA but more slowly than plaque-forming ability.

**Carcass removal.** The carcass removal rate (λ*_D_* = 0.7 day^-1^) is based on work showing rapid scavenging and decay of carcasses (Harp and Petranka 2006, Le Sage et al. 2019).

**Low-intensity infection recovery.** The recovery pathway γ is set to zero in the baseline parameterization, because recovery from low-intensity ranavirus infection is not well-documented in larval amphibians. The effect of nonzero γ is evaluated in the sensitivity analysis (see main text; Appendix S1: Figure S5).

**Model variant–specific parameters.** For Model A, we assigned a shedding rate equal to the geometric mean of the low- and high-intensity values (√(ρ_low_ · ρ_high_) = 5,000 copies day^-1^), the midpoint on the log scale across which shedding rates span, to avoid biasing the estimate toward the much larger high-intensity value. For Model B, the constant progression rate (0.125 day^-1^) equals σ(*W*) evaluated at the half-saturation concentration (*W* = *K*_σ_), providing a comparison that removes dose-dependence while fixing progression at the half-maximum exposure-dependent rate.

### Section S3: Extended Results

**Environmental-transmission Hill coefficient:** At the baseline shedding ratio (10^4^), both *n* = 1 and *n* = 2 produce >99.8% cumulative mortality and qualitatively similar dynamics. The sharp mortality threshold remains under both parameterizations, with the 50% mortality boundary occurring at nearly the same shedding ratios (Figure S3). Environmental transmission contributes to the disease-free-equilibrium linearization when *n* = 1 but not when *n* = 2. *R*_0_ is undefined under the baseline assumption σ_0_ = 0 and γ = 0 for both coefficients because the transition matrix **V** is singular.

**Progression function form**: Hill vs. piecewise-linear progression functions produce cumulative mortality within 3 percentage points of each other at baseline parameters. The step-function formulation (σ = 0 for *W* < *K*_σ_, σ = σ_max_ for *W* ≥ *K*_σ_) fails to produce die-offs when σ_0_ = 0 because no high-intensity infections are generated until *W* reaches *K*_σ_, but low-intensity shedding by itself cannot raise *W* to *K*_σ_ (maximum low-intensity *W* ≈ 0.40 copies/mL << *K*_σ_ = 25 copies/mL).

**Transmission form**: Frequency-dependent (β*_I_* · *I*/*N*) vs. density-dependent (β*_I_* · *I*, with β*_I_* rescaled to match the initial force of infection) contact transmission produces qualitatively identical dynamics (mortality within 0.01 percentage points). The frequency-dependent formulation makes *R*_0_ independent of density and volume, simplifying interpretation.

**Carcass contribution**: Removing carcass leaching (ρ*_D_* = 0) reduces peak *W* from 1,790 to 1,499 copies/mL (16% reduction) but does not eliminate die-offs (cumulative mortality 99.9%). Carcasses contribute to viral persistence but are not required for the modeled environmental feedback.

**Extended sensitivity results.** Extended sensitivity analysis across wider parameter ranges identifies sharp changes in cumulative mortality for each of the six parameters spanning the full mortality range in the one-at-a-time analysis. Die-offs fail to occur when σ_max_ falls below ~15% of its default value, when *K*_σ_ exceeds 3× default, or when pond volume exceeds 3× default with the baseline host population size. Cumulative mortality exceeds 97% for δ = 0.1–0.6 day^-1^ (Figure S7), spanning from slow eDNA degradation to moderately rapid viral inactivation under filter-sterilized conditions (Johnson & Brunner 2014). At higher decay rates (δ ≥ 1.0 day^-1^), the environmental reservoir clears too rapidly for viral accumulation to raise *W* to concentrations at which σ(*W*) increases rapidly at the baseline *K*_σ_, though reducing *K*_σ_ restores die-off dynamics even at δ = 2.0 (Figure S8). A rescaling invariance test shows that proportionally rescaling all concentration-dependent parameters (ρ_high_, ρ_low_, ρ_D_, *K*_σ_, and *K*_inf_) across four orders of magnitude leaves cumulative mortality and onset timing unchanged (Figure S9), showing that the model's behavior depends on ratios among these parameters rather than their absolute values.

| **Table S1.** Basic reproduction number (*R*_0_) and cumulative mortality for each model variant at baseline parameters (*N*_0_ = 20,000, *n* = 2). | | | | |
| --- | --- | --- | --- | --- |
| **Model** | ***R*_0_** | **Cumulative mortality (%)** | **Time to 50% mortality (days)** | **Interpretation** |
| A: SIWR | 1.4 | 95.4 | 58.1 | Single infected class; high mortality via typical epizootic (no sharp mortality threshold) |
| B: Constant-progression | 3.0 | 100 | 23.5 | Staging + asymmetry produce rapid mortality |
| C: Dose-dependent | undefined | 99.9 | 45.8 | Contact transmission permits invasion; environmental viral accumulation drives dose-dependent progression and die-off |

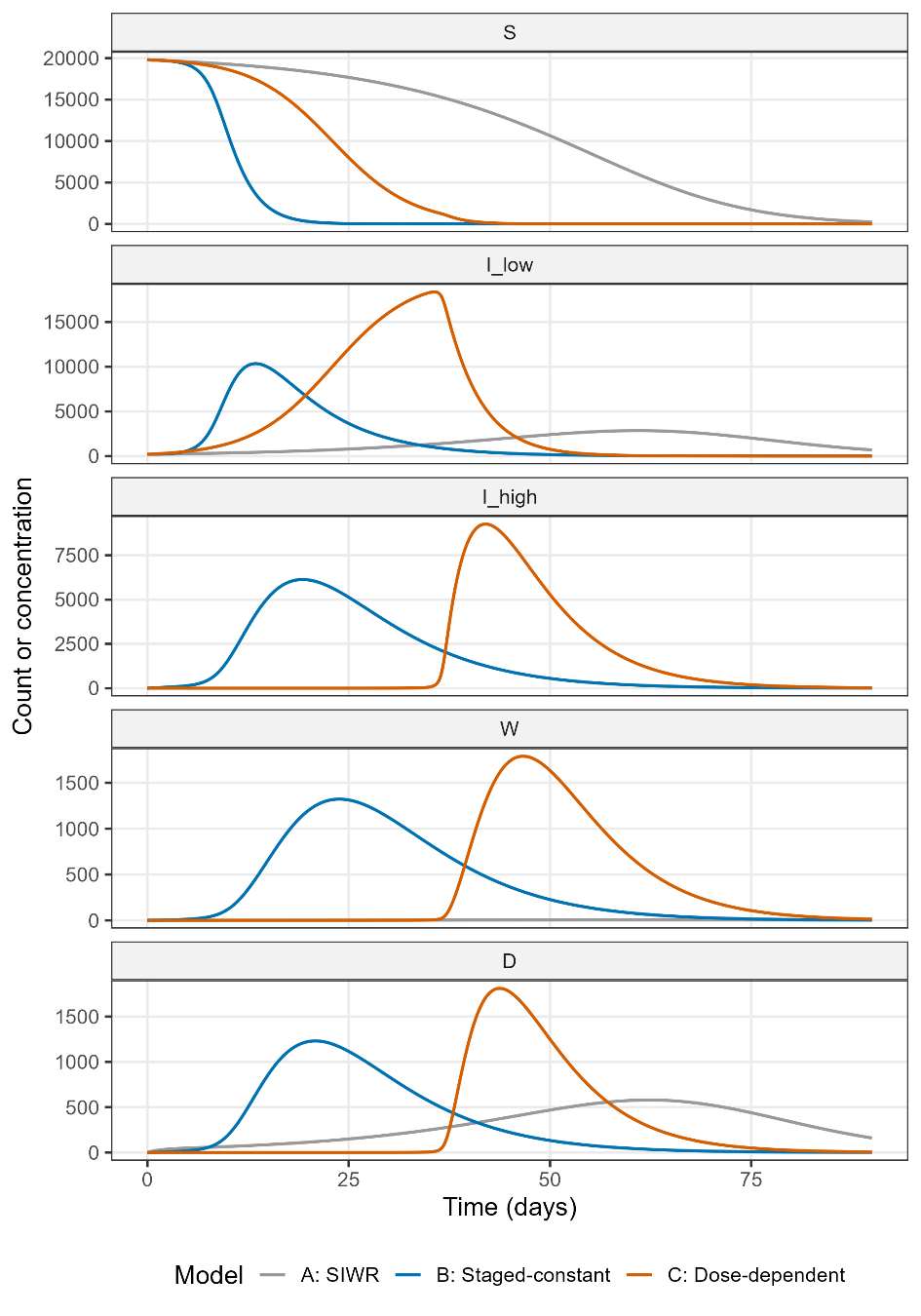

**Figure S1. Baseline dynamics for all compartments across three model variants.** Time series of all five state variables (susceptible hosts (*S*), low-intensity infections (*I*_low_), high-intensity infections (*I*_high_), environmental virus (*W*, copies/mL), and carcasses (*D*)) for Model A (SIWR, grey), Model B (staged-constant, blue), and Model C (dose-dependent, orange) at default parameters (*N*_0_ = 20,000, *V* = 10,000 L). For Model A, the single infected compartment is shown in the *I*_low_ panel; Model A has no separate high-intensity infection compartment.

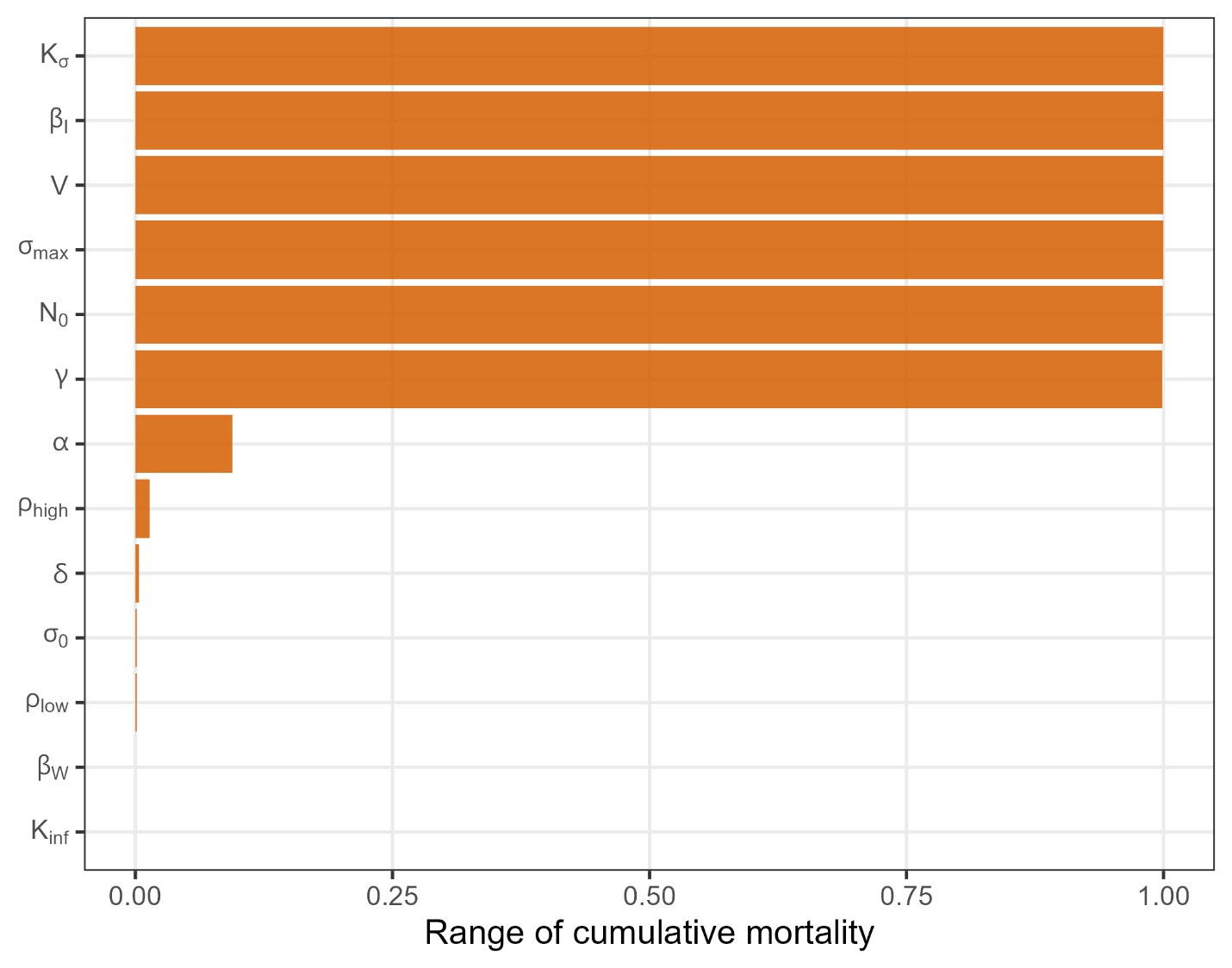

**Figure S2. One-at-a-time parameter sensitivity.** Range of cumulative mortality when each parameter is varied individually across its plausible range (Table 1) while holding all others at default values. Six parameters (σ_max_, β*_I_*, *K*_σ_, *V*, *N*_0_, γ) span the full or nearly full 0–100% mortality range; the low-intensity shedding rate (ρ_low_) and disease-induced mortality rate (α) have negligible individual influence (range < 0.094).

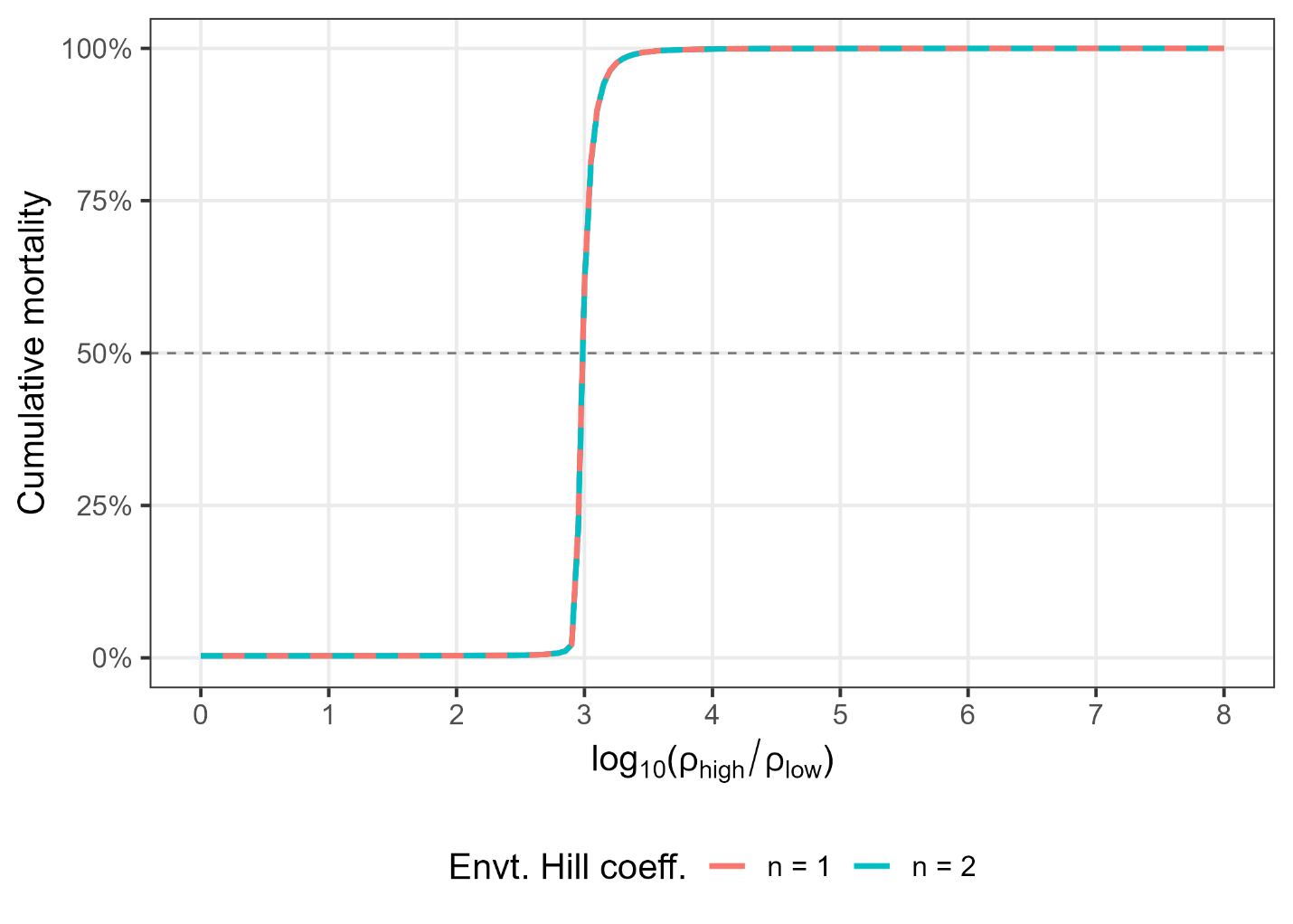

**Figure S3. Robustness to environmental transmission Hill coefficient.** Cumulative mortality across the high-to-low shedding-ratio range for *n* = 1 (Michaelis-Menten) and *n* = 2 (sigmoidal).

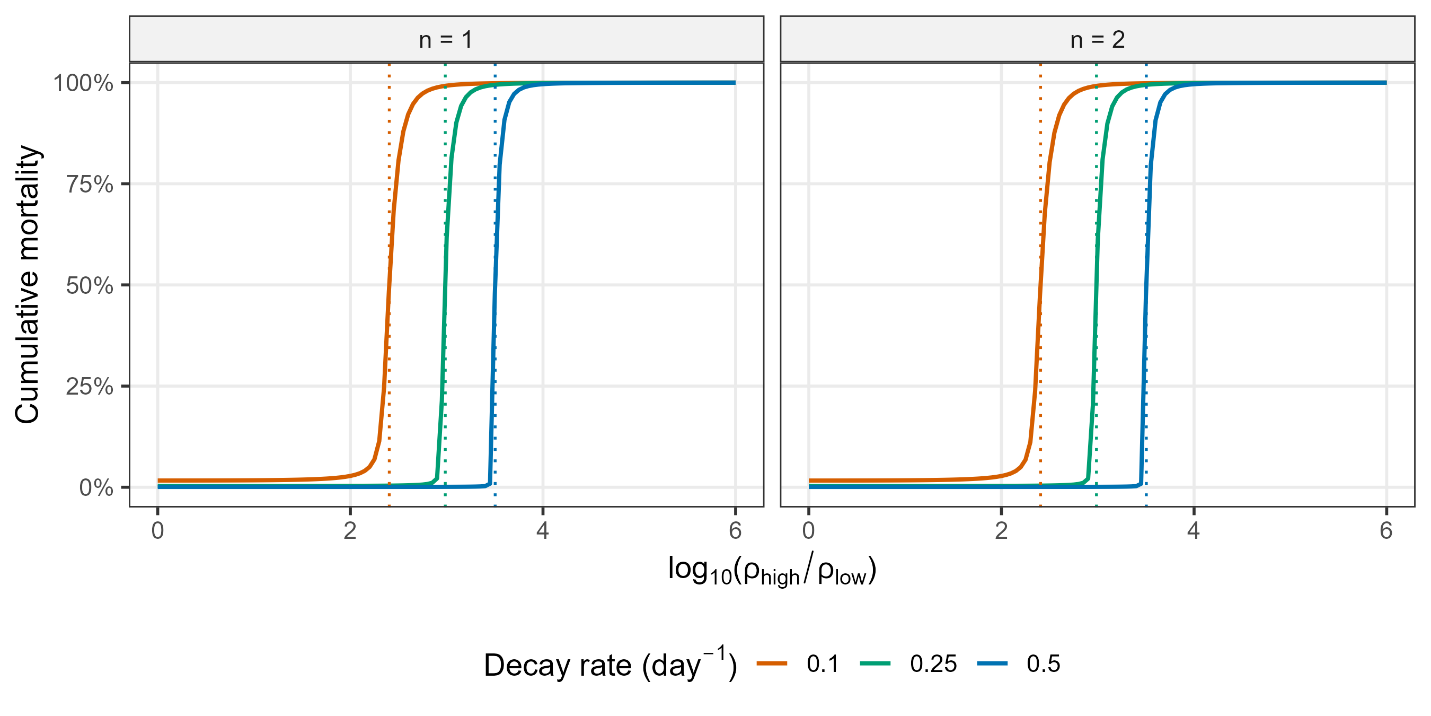

**Figure S4. Structural robustness: Hill coefficient × viral decay rate.** Cumulative mortality across the high-to-low shedding-ratio range for *n* = 1 (left) and *n* = 2 (right) across three viral decay rates (δ = 0.1, 0.25, 0.5 day^-1^). Dotted vertical lines mark the shedding ratio associated with 50% cumulative mortality for each decay rate.

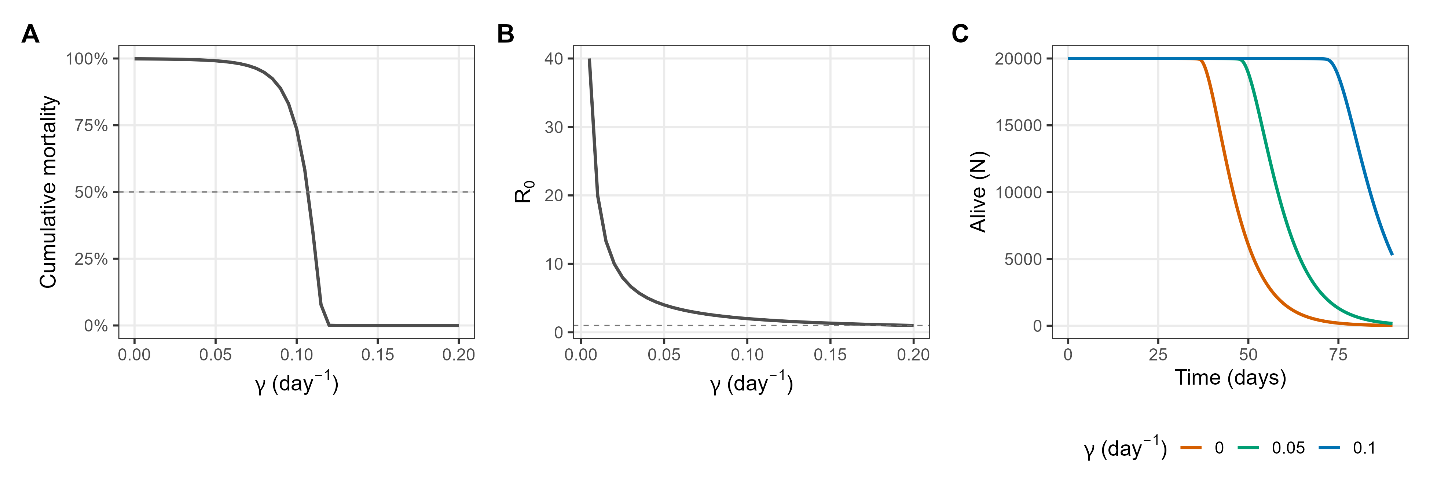

**Figure S5. Sensitivity to low-intensity infection recovery rate (γ).** (A) Cumulative mortality as a function of recovery rate. (B) *R*_0_ as a function of γ; dashed line marks *R*_0_ = 1. (C) Population dynamics (total alive, *N*) at three recovery rates: γ = 0 (baseline, orange), γ = 0.05 (green), and γ = 0.1 day^-1^ (blue).

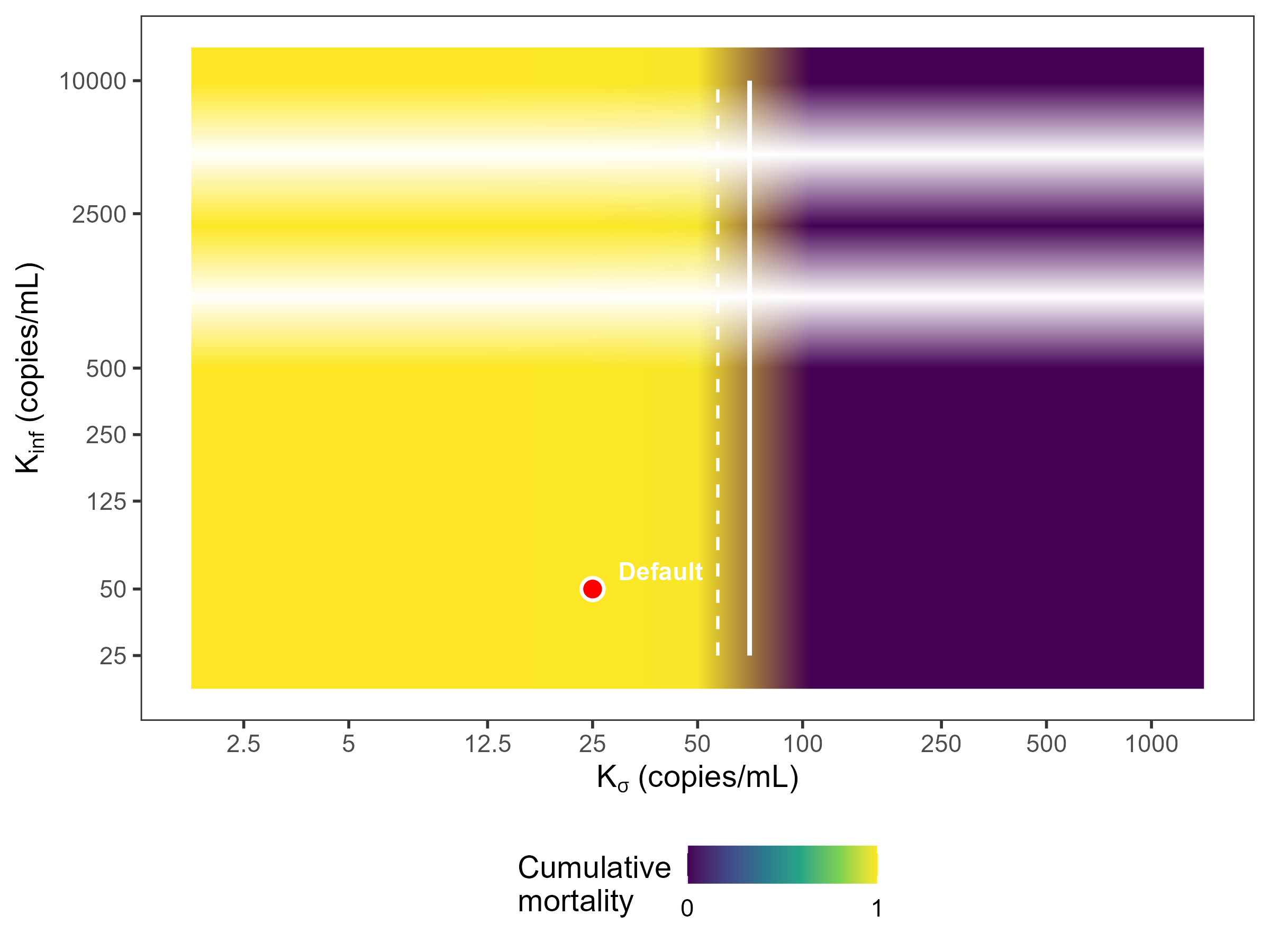

**Figure S6. Joint sensitivity of half-saturation constants *K*_σ_ and *K*_inf_.** Heatmap of cumulative mortality across the *K*_σ_ (progression half-saturation) and *K*_inf_ (infection half-saturation) parameter space. Solid and dashed contours mark 50% and 80% cumulative mortality, respectively. Red dot marks the default setting (*K*_σ_ = 25, *K*_inf_ = 50 copies/mL).

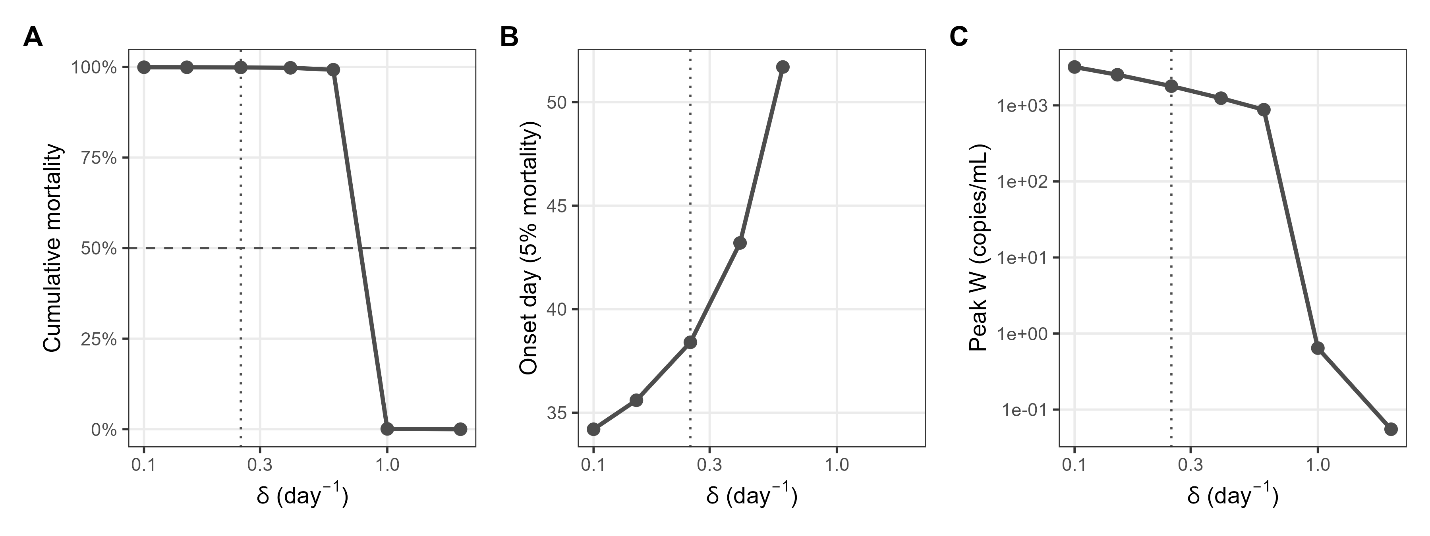

**Figure S7. Sensitivity to viral decay rate (δ).** (A) Cumulative mortality, (B) die-off onset timing (day of 5% cumulative mortality), and (C) peak environmental viral concentration (log_10_ scale) as functions of δ. Dotted vertical line marks the default value (δ = 0.25 day^-1^, half-life ≈ 2.8 days).

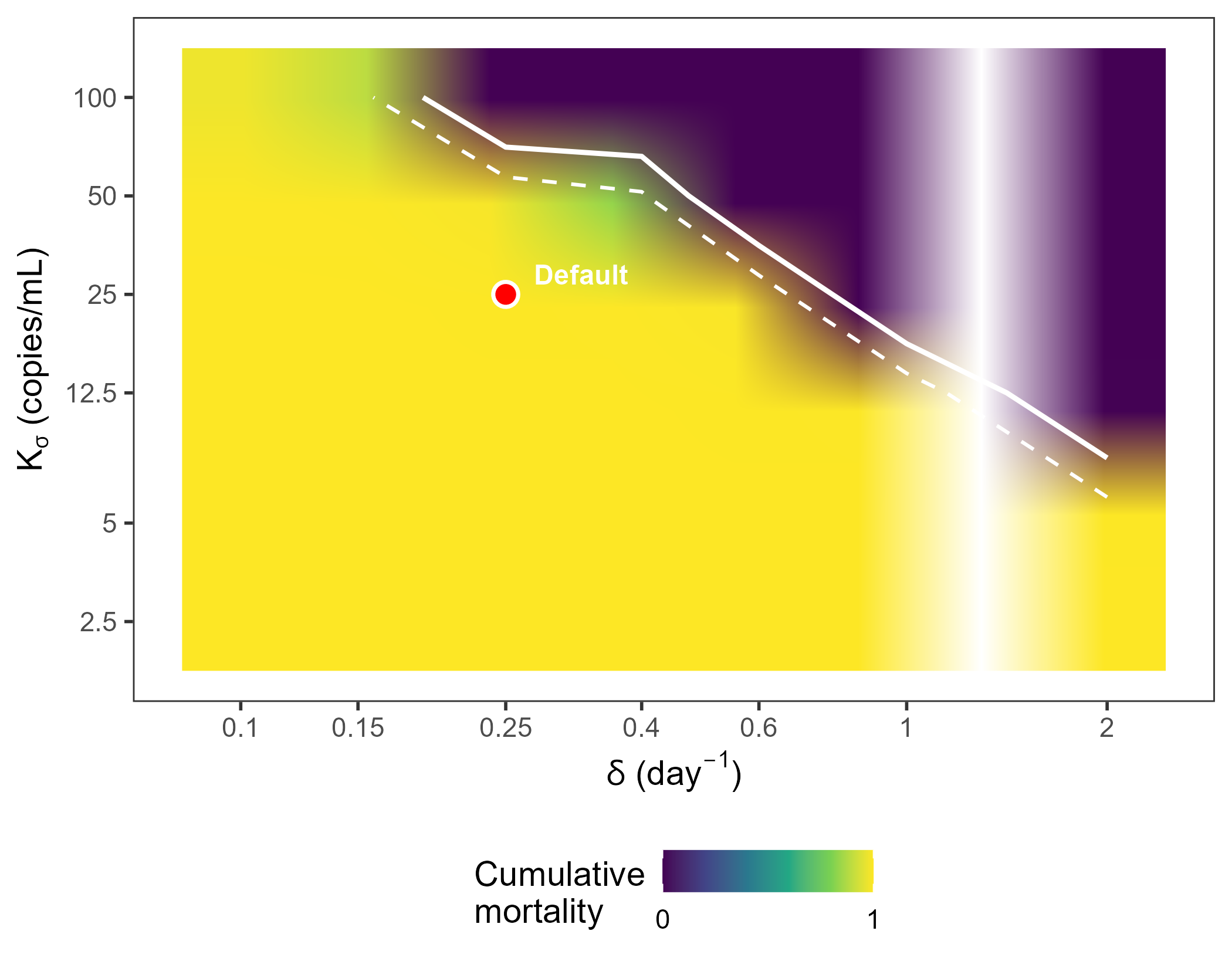

**Figure S8. Joint sensitivity of viral decay rate and progression half-saturation concentration.** Heatmap of cumulative mortality across the δ × *K*_σ_ parameter space. Solid and dashed contours mark 50% and 80% cumulative mortality, respectively. Red dot marks the default operating point (δ = 0.25 day^-1^, *K*_σ_ = 25 copies/mL).

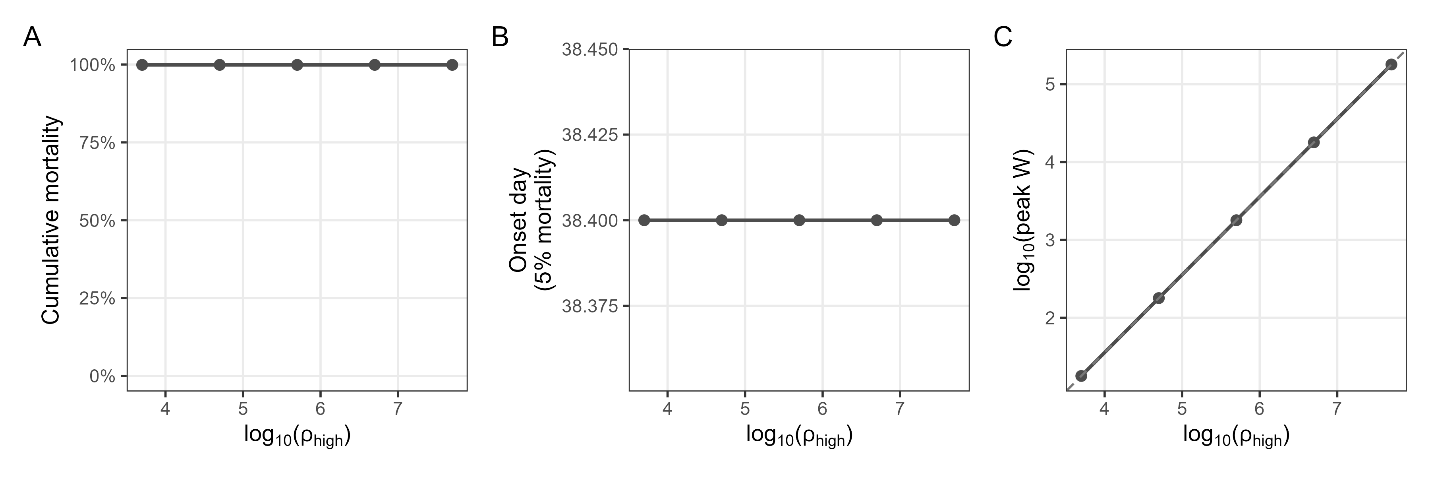

**Figure S9. Rescaling invariance of shedding and half-saturation parameters.** (A) Cumulative mortality, (B) onset timing, and (C) peak *W* when ρ_high_, ρ_low_, ρ_D_, *K*_σ_, and *K*_inf_ are proportionally rescaled across four orders of magnitude while maintaining their ratios.

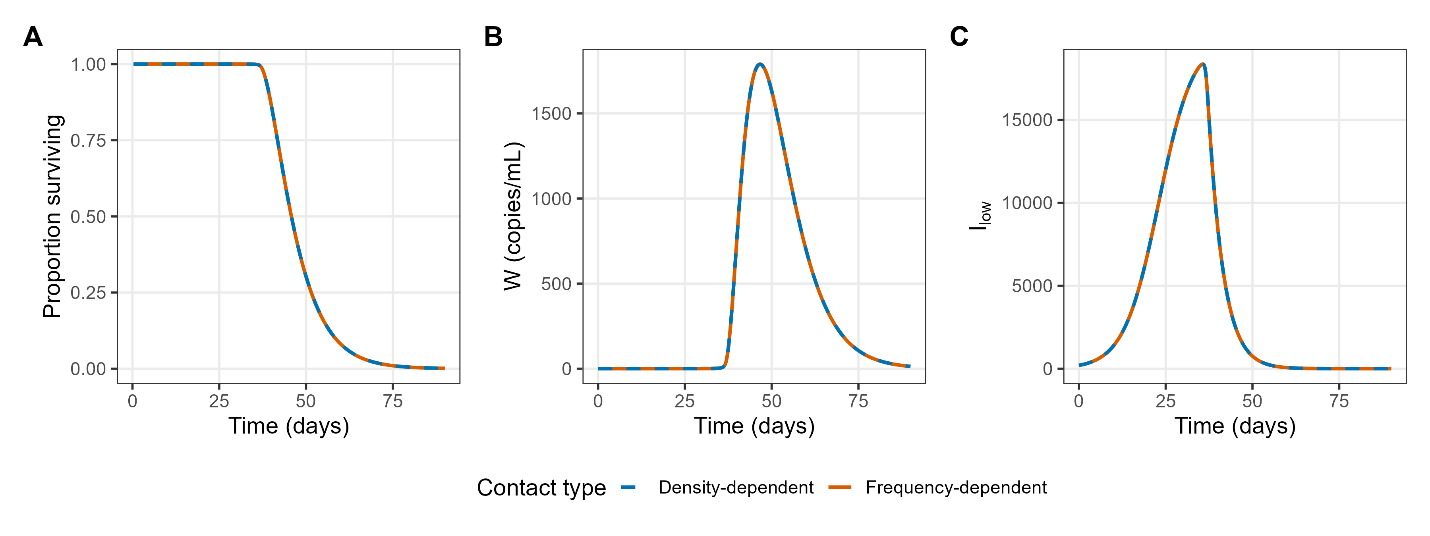

**Figure S10. Frequency-dependent vs. density-dependent contact transmission**. Comparison of Model C dynamics under frequency-dependent (β*_I_* · *I*/*N*; solid orange) and density-dependent (β*_I_* · *I*; dashed blue, with β*_I_* rescaled to match initial force of infection) contact transmission at baseline parameters. (A) Proportion of population surviving, (B) environmental viral concentration (copies/mL), and (C) low-intensity infections.

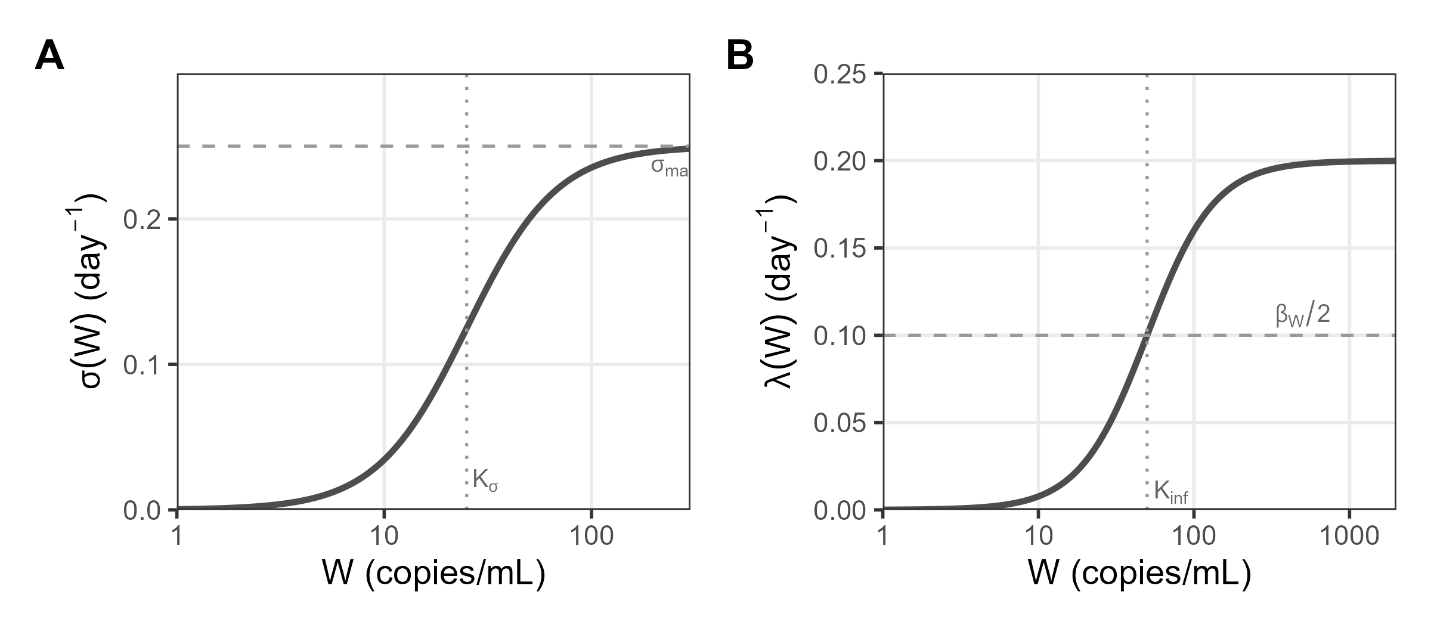

**Figure S11.** **Dose-response functions governing environmental transmission and infection progression.** (A) The Hill-type progression function σ(*W*) (*K*_σ_ = 25 copies/mL, σ_max_ = 0.25 day^-1^), which governs the rate of progression from low- to high-intensity infection as a function of environmental viral concentration *W*. Dashed horizontal line marks σ_max_; dotted vertical line marks *K*_σ_. (B) The saturating environmental force of infection λ(*W*) (*K*_inf_ = 50 copies/mL, β*_W_* = 0.2 day^-1^), which governs the rate of new infections from the environmental reservoir. Dashed horizontal line marks β*_W_*/2; dotted vertical line marks *K*_inf_. The environmental-transmission and progression functions use Hill coefficients *n* = 2 and *m* = 2.

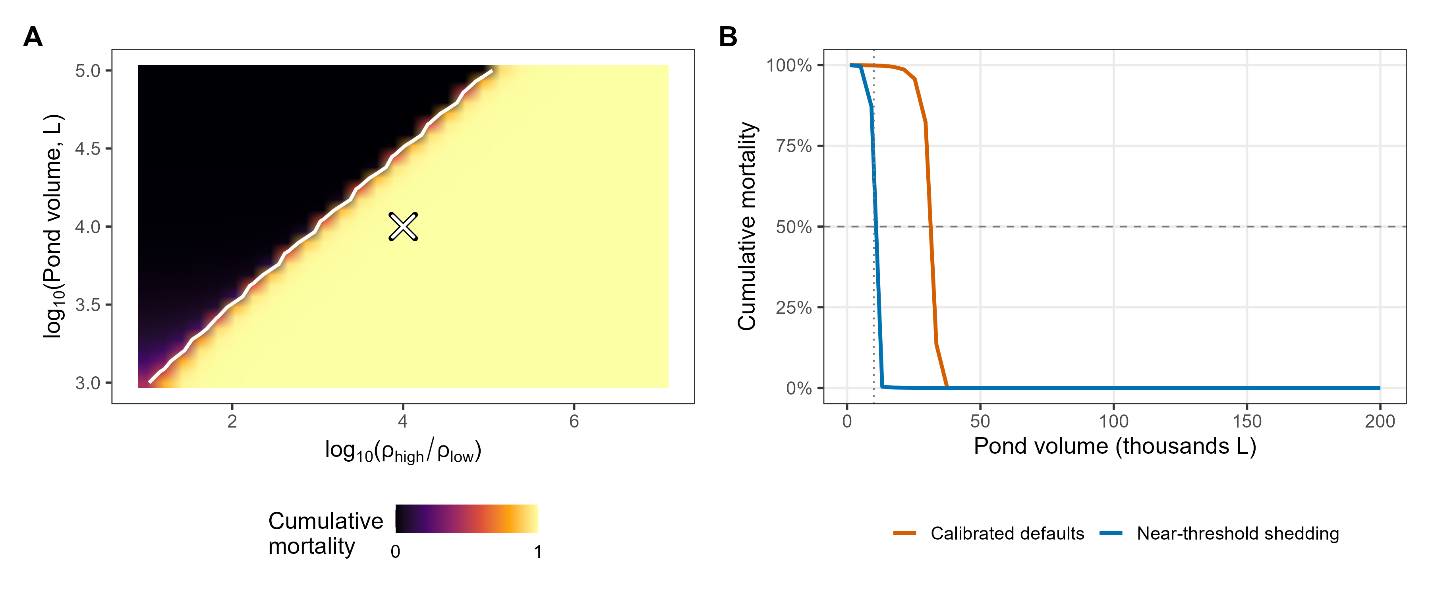

**Figure S12.** Joint shedding-ratio and volume sensitivity. (A) Cumulative mortality across the high-to-low shedding ratio (log_10_(ρ_high_/ρ_low_)) and pond volume (log_10_(*V*); inferno color scale) at fixed *N*_0_  = 20,000. The white contour marks 50% mortality; the cross marks the baseline parameterization (ρ_high_/ρ_low_ = 10^4^, *V* = 10^4^ L). (B) Cumulative mortality as a function of pond volume at constant host number (*N*_0_ = 20,000), for calibrated defaults (ρ_high_ = 500,000; orange) and near-threshold shedding (ρ_high_ = 50,000; ρ_high_/ρ_low_ = 10^3^; blue). Larger ponds dilute environmental virus and can prevent *W* from reaching concentrations at which σ(*W*) increases rapidly. The dotted vertical line marks the default volume (10,000 L); the dashed horizontal line marks the 50% die-off threshold.
